# *SMPD4* mediated sphingolipid metabolism regulates brain and primary cilia development

**DOI:** 10.1101/2023.12.15.571873

**Authors:** Katherine A. Inskeep, Bryan Crase, Rolf W. Stottmann

## Abstract

Genetic variants in multiple sphingolipid biosynthesis genes cause human brain disorders. A recent study collected patients from twelve unrelated families with variants in the gene *SMPD4*, a neutral sphingomyelinase which metabolizes sphingomyelin into ceramide at an early stage of the biosynthesis pathway. These patients have severe developmental brain malformations including microcephaly and cerebellar hypoplasia. However, the mechanism of *SMPD4* was not known and we pursued a new mouse model. We hypothesized that the role of *SMPD4* in producing ceramide is important for making primary cilia, a crucial organelle mediating cellular signaling. We found that the mouse model has cerebellar hypoplasia due to failure of Purkinje cell development. Human induced pluripotent stem cells exhibit neural progenitor cell death and have shortened primary cilia which is rescued by adding exogenous ceramide. *SMPD4* production of ceramide is crucial for human brain development.

## Introduction

The human brain is the most unique and complicated structure in the body. Neurological disorders represent a significant healthcare burden and research emphasis. The ability to sequence the human genome has greatly facilitated the study of genetic variants causing rare monogenic pediatric brain disorders. This model was recently illustrated by the publication of a cohort of 23 patients from 12 unrelated families with variants in *SMPD4* (Magini et al., 2019). These patients share common phenotypes of congenital microcephaly (primary microcephaly 15/21, progressive microcephaly 9/10), cerebellar hypoplasia (10/21), delayed or hypomyelination of the brain, other structural brain abnormalities, and severe developmental delay (7/7). Patients also commonly presented with intrauterine growth restriction, neonatal respiratory distress, congenital arthrogryposis of hands and feet, and abnormal muscle tone (Magini *et al*., 2019). Two more pediatric patients with similar phenotypes have since been described (Bijarnia-Mahay et al., 2022; Yamada et al., 2022) along with three surviving adult patients from one family who present with microcephaly, ataxia, severe developmental disability, and insulin-dependent diabetes (Smits et al., 2023).

Microcephaly is defined by a head circumference less than three standard deviations below the mean for age and sex, resulting in intellectual disability and developmental delay (Dobyns, 2002; Woods et al., 2005). In the United States, 1:800-5,000 babies are born with microcephaly annually; or about 25,000 US births per year (Becerra-Solano et al., 2021). The development of the human cerebral cortex is a complex process with multiple steps including proliferation and differentiation of progenitor cells, migration of neuronal precursors into the cortical plate, and organization into the six layers of the mature cortex (Lee, 2019; Lee and 2019; Molnár et al., 2012; Molnár et al., 2019). The balance between proliferative and asymmetric divisions of progenitor cells in the ventricular zone is essential for generating the human brain’s complex and expanded cerebral cortex relative to other species. Many of the genes tied to primary microcephaly in humans are associated with the centrosome and mitotic spindle, which regulate cell division (Bond et al., 2002; Bond et al., 2005; Gilmore and Walsh, 2013; Guernsey et al., 2010; Jackson et al., 2002; Kumar et al., 2009). Similarly, defects in primary cilia-associated genes cause cell cycle and neural progenitor defects (Gilmore and Walsh, 2013; Jayaraman et al., 2016; Zhang et al., 2019). While there are species-specific differences between mouse and human corticogenesis, mouse models have generally served as a reliable model for these facets of cortical development (Capecchi and Pozner, 2015; Gruber et al., 2011; McIntyre et al., 2012; Putkey et al., 2002).

Cerebellar hypoplasia is characterized simply by smaller cerebellar size and global underdevelopment (Accogli et al., 2021). Cerebellar granule cells are the most abundant cell type in the mammalian brain. Unlike in the cerebral cortex, the cerebellar germinal zone is closest to the pial surface, and granule cells migrate internally after birth as the cerebellum develops (Altman and Bayer, 1985a; Fujita, 1967). The *Sonic hedgehog* (SHH) pathway is critical for cerebellar development (Wechsler-Reya and Scott, 1999). Purkinje cells are the only output neurons of the cerebellar cortex and are not produced from the external granule layer (EGL) but do secrete SHH required for granule cell proliferation (Butts et al., 2014; Corrales et al., 2004; Fujita, 1967; Seto et al., 2014). Granule cell progenitors (GCPs) and Purkinje cells are ciliated (Del Cerro and Snider, 1972; Doughty et al., 1998) and these primary cilia are required for their survival, integrity, and localization (Bowie and Goetz, 2020). Accordingly, many ciliopathy disorders affect the cerebellum (Abdelhamed et al., 2013; Epting et al., 2020; Inskeep et al., 2022; Lee and Gleeson, 2011). Cortical and cerebellar development have both similarities and intriguing differences that become relevant in our study.

Neutral sphingomyelinase 3 (*SMPD4*) is a member of the sphingomyelinase family, which hydrolyzes sphingomyelin to produce ceramide and phosphorylcholine in the catabolic pathway of sphingolipid biosynthesis (Krut et al., 2006). Ceramide is the precursor of most complex sphingolipids. Disorders of sphingolipid content include Krabbe’s disease, Gaucher’s disease, and Niemann-Pick disease (*SMPD1*) (Platt, 2014). These lipid storage disorders all involve improper accumulation of sphingolipids or lipid components in the lysosome. Moreover, lipid rafts formed from sphingolipids are involved in dementia, Parkinson’s, and Alzheimer’s disease (Mencarelli and Martinez-Martinez, 2013). Ceramide is a cornerstone lipid with diverse roles in the cell. Studies have linked cell stress and apoptosis induction to increased ceramide levels (Herget et al., 2000; Kolesnick and Krönke, 2003; Lee et al., 2015). However, its roles in cell division and primary ciliogenesis are of particular interest to our study. Ceramide interacts with acetylated tubulin to stabilize centrosome-associated microtubules (Tripathi et al., 2021). It is well known that ceramide is absolutely required for primary ciliogenesis, and specifically in neural progenitor cells (He et al., 2014; Wang et al., 2009). Primary cilia and centrosome cycling are inextricably linked to the cell cycle (Kim and Tsiokas, 2011).

In this study, we sought to establish a mouse model and study the molecular mechanism of *SMPD4* mediated structural brain malformations. First, we determined the developmental timeline of *Smpd4* expression in mouse brain. We found *Smpd4* is highly expressed in developing forebrain, particularly in the ventricular zone, starting at embryonic day (E)12.5, and in both granule and Purkinje cells in the cerebellum from E14.5. We present an *Smpd4* null mouse model with incompletely penetrant perinatal lethality, failure to thrive, and cerebellar hypoplasia. We further determined the cause of cerebellar defect is likely due to loss of Purkinje cells. *Smpd4* null and forebrain-specific knockout mice do not exhibit the microcephaly seen in human patients. We established human stem cell models which showed a loss of neural progenitor cells due to cell death and decreased proliferation. We explored the role of primary cilia in these models and found that although *Smpd4* null mice appear unaffected, human induced pluripotent stem cell (iPSCs) models have shortened and bulbous primary cilia. We propose that this is due to changes in subcellular ceramide, as treating iPSCs with exogenous ceramide significantly rescues cilia length. We also found evidence for disrupted WNT pathway signaling. We suggest that during brain development, *SMPD4* provides ceramide in an organelle-specific manner, regulating the cell cycle and allowing primary cilia to grow properly.

## Results

### Smpd4 is highly expressed in the developing mouse brain

A comprehensive analysis of *Smpd4* expression in the developing mouse brain has not been previously carried out. Human *SMPD4* is expressed in most tissue types throughout embryonic and postnatal development including brain and human-derived neural organoids (Fietz et al., 2012; Luo et al., 2016; Zhu et al., 2016). Similar RNA sequencing experiments in mouse show *Smpd4* expression in multiple brain regions including the ventricular zone (VZ), subventricular zone (SVZ), and cortical plate (Fietz *et al*., 2012). However, these studies do not detail *Smpd4* expression in the developing forebrain or cerebellum during development.

Therefore, we assayed mRNA forebrain expression via RNAScope probes at E12.5 through P27. We first noted *Smpd4* expression in the forebrain at E12.5 (data not shown). Expression is highest at E14.5, the peak of neurogenesis in the mouse cortex corresponding to the end of the second trimester in human development (Chen et al., 2017) (**Fig 1A**). We analyzed serial sections of E14.5 and E18.5 mouse cortex stained with *Smpd4* and PAX6 to confirm that *Smpd4* is expressed in the VZ, overlapping neural progenitor cells (Estivill-Torrus et al., 2002; Heins et al., 2002) (**Fig 1D-F, Supp Fig 1A-C**). We found that *Smpd4* is gradually restricted to the VZ by E18.5 (**Fig 1B**) but is maintained at low levels across the cortex until adulthood (**Fig 1C**). Our data show *Smpd4* is expressed in the developing cortex at critical stages and cell types for neurodevelopment.

**Figure 1:**
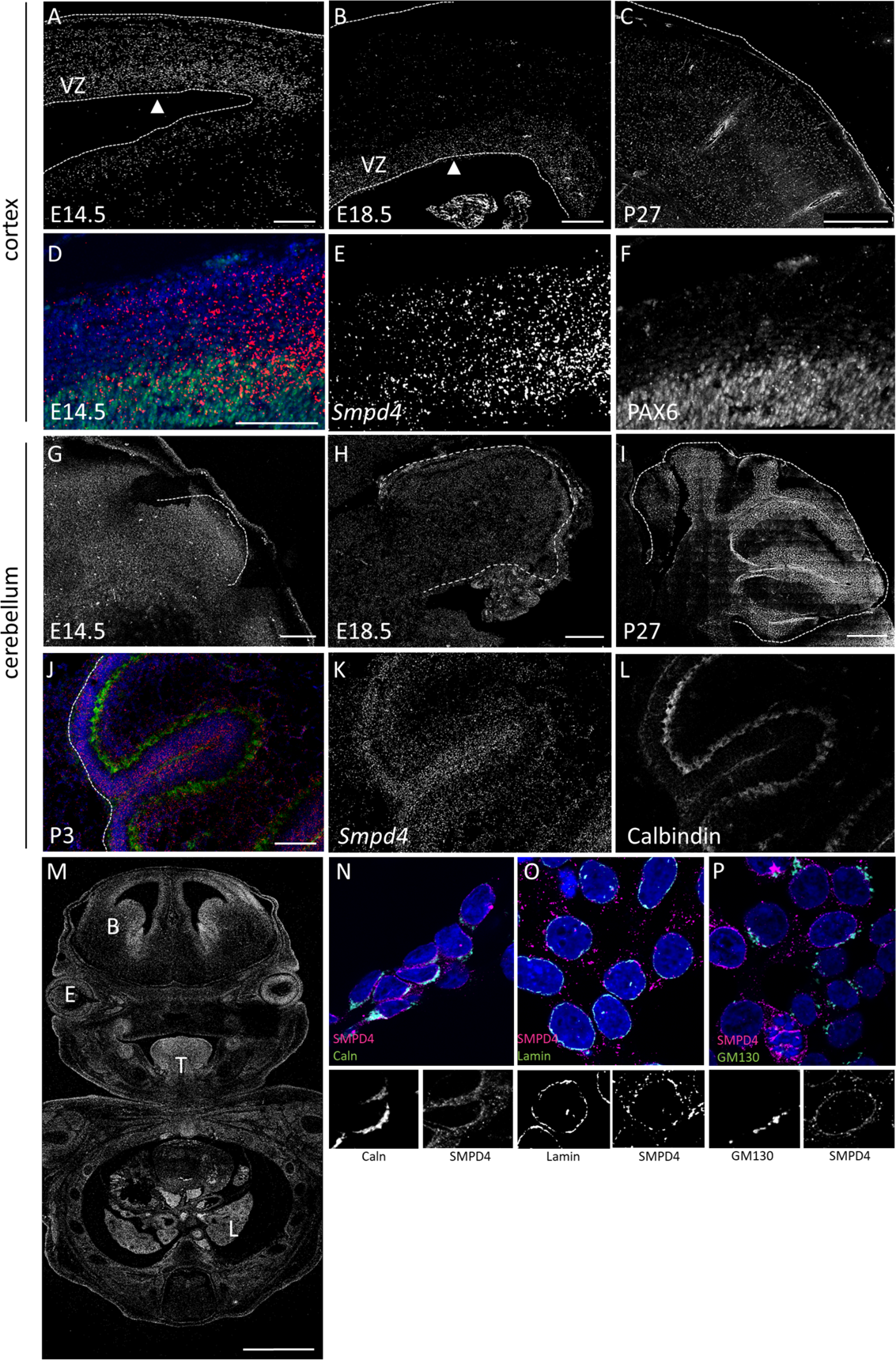
*Smpd4* is expressed throughout mouse cortical and cerebellar development. *Smpd4* expression in mouse cortex at E14.5, E18.5, and P27. Ventricular surface is marked with an arrowhead, VZ= ventricular zone (A-C). Serial sections of *Smpd4* RNA expression and PAX6 immunohistochemistry show that *Smpd4* colocalizes with ventricular zone progenitor cells but further extends across the cortex at E14.5 (D-F). *Smpd4* expression in mouse cerebellar primordium at E14.5 (G) and E18.5 (H) and postnatal cerebellum (P27, I). In the P3 cerebellum, *Smpd4* is expressed in both the external granule cell layer and Calbindin-positive Purkinje cells, (J-L). Analysis of the whole embryo at E14.5 shows *Smpd4* expression in the brain, eyes, tongue, and lungs (M, B=brain, E=eye, T=tongue, L=Lung). In HEK293T cells, SMPD4 does not colocalize with GM130 in the Golgi apparatus (N) but is found in the Calnexin-positive ER (O) and Lamin-positive nuclear membrane (P). Scale bars in A,B,D,J =100μm C,I=500μm, G,H= 200μm, M=1000μm.

Additionally, we observed widespread *Smpd4* expression at very early stages in the cerebellar primordium at E14.5 (**Fig 1G**). Expression is maintained to maturation at P21 (**Fig 1H,I**). *Smpd4* is expressed in multiple cell types of the cerebellum, most highly in granule cells but also in

Calbindin-positive Purkinje cells (**Fig 1J-L, Supp Fig 1D-F**). In the developing mouse embryo at E14.5, *Smpd4* is widely expressed but especially in the brain, lungs, tongue, and eye (**Fig 1M**). Further, we assessed subcellular localization of SMPD4 in human HEK293T cells. At the level of the organelle, SMPD4 localizes to both the Calnexin-positive endoplasmic reticulum (**Fig 1N**) and Lamin-positive nuclear membrane (**Fig 1O**), but not the GM130-positive Golgi apparatus (**Fig 1P**), unlike previously reported (Stoffel et al., 2016). We have detailed the expression of *Smpd4* in neurogenesis and found it is consistent with a role in both cortical and cerebellar development.

### Loss of Smpd4 causes perinatal demise, failure to thrive, and cerebellar hypoplasia

Mouse models serve as a tool to understand pathogenesis of human disease. We hypothesized that loss of *Smpd4* in a null mouse model would lead to structural brain malformations consistent with our expression analysis and a previous human cohort. The International Mouse Phenotyping Consortium generated a conditional gene trap allele of *Smpd4* (*Smpd4^tm2a(KOMP)Wtsi^*). A null allele generated from these mice, *Smpd4^tm2b^* (i.e., *Smpd4^null^*), was observed to have incompletely penetrant lethality at weaning but noted no further developmental abnormalities. However, this analysis was limited to only surviving adult animals. We used this line to determine the role of *Smpd4* in embryonic and postnatal brain development.

In our hands, a significant majority of *Smpd4^null/null^* mice die prior to P0 (n=6 survivors of 10.75 expected, p=0.15, **Fig 2A**). Only 4% survived to weaning at P21 (n=7/23 expected, p<0.001, **Fig 2B, Supp Table 4**). Surviving animals all failed to thrive, were about half the weight of their littermates (n=6, p<0.001, **Fig 2C,D**), had no sex bias, and subsequently required euthanasia. *Smpd4^null/null^* animals had significantly smaller brains as measured by weight (**Fig 2E**) and dorsal surface area at P0 (n=4, p<0.001, **Fig 2F,G,L**) and P21 (**Fig1 H-K)**. We were unable to determine a specific molecular cause for the reduced brain size (**Supp Fig 2**). The overall body size in mutants is also significantly smaller at P21 (**Fig 2D**). Although the human definition of microcephaly does not account for body size (Gilmore and Walsh, 2013), the *Smpd4* mouse model does not show a disproportionately small forebrain in surviving mutants.

**Figure 2:**
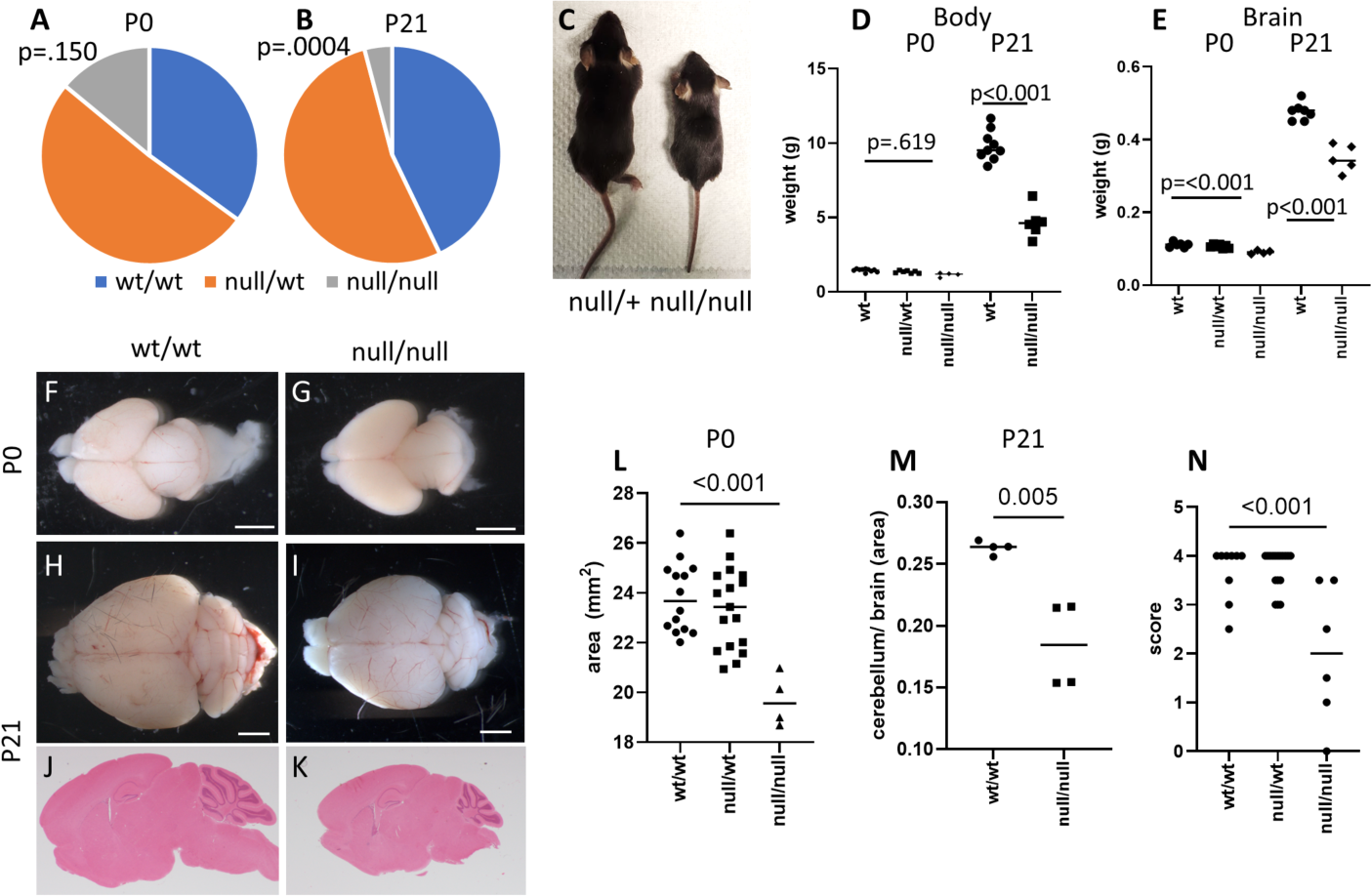
Loss of *Smpd4* causes perinatal demise, cerebellar hypoplasia and microcephaly in surviving animals. *Smpd4* homozygous null animals survive at sub-Mendelian ratios after birth (A,B). They are significantly smaller than their littermates at weaning but not at birth (C,D) and have smaller brains at all stages examined (E). Homozygous null brains are mostly structurally intact at P0 (F,G) and P21 (H-K), though they exhibit some microcephaly (L) and cerebellar hypoplasia (M). At P21, homozygous null animals exhibit hindlimb clasping indicating a cerebellar deficit (N).

Conversely, we were particularly intrigued that the surviving *Smpd4^null/null^* mice have severe cerebellar hypoplasia even after controlling for smaller brain size (**Fig 2M**). At weaning, both males and females demonstrated hindlimb clasping consistent with a cerebellar deficit (**Fig 2N**). There was no statistical difference in body or brain weight between P21 males and females (p=0.991). Overall, these brain and behavior phenotypes are partially consistent with those observed in published human *SMPD4* patients (Magini *et al*., 2019).

### Purkinje cells fail to support cerebellar postnatal development in the absence of Smpd4

To circumvent the apparent early postnatal lethality of the *Smpd4* germline model, we decided to employ a conditional genetic strategy. We utilized the Cre-lox system, in which loxP sites surrounding a critical exon in a gene of interest (i.e. ‘flox’) can be combined with tissue-specific Cre recombinase expressing transgenic mice. This affords specific spatiotemporal control in deleting the gene of interest. We used Cre transgenic lines to specifically delete *Smpd4* in three discrete lineages: the hindbrain (*Engrailed1-*Cre), dorsal telencephalon (*Emx1-*Cre), and forebrain (*Foxg1-*Cre).

Motivated by cerebellar hypoplasia in *SMPD4* patients and the *Smpd4* germline model, we generated a hindbrain-specific deletion of *Smpd4* using *Engrailed1*-Cre (*En1*-Cre). *En1-Cre+; Smpd4^flox/null^* animals survived to weaning (**Supp Table 5**) but were smaller and ataxic as compared to their littermates.

While the cerebellum was morphologically normal in early postnatal cerebellar development (P5, **Fig. 3A**), it appeared obviously smaller at P14 (**Fig 3B**). P21 *En1-Cre+; Smpd4^flox/null^* animals have readily apparent cerebellar hypoplasia (**Fig 3C,D, quantified in E**). Specifically, they have reduced vermian height, hemisphere width, and overall cerebellar width, without reduced vermian width (**Fig 3G-J**).

**Figure 3:**
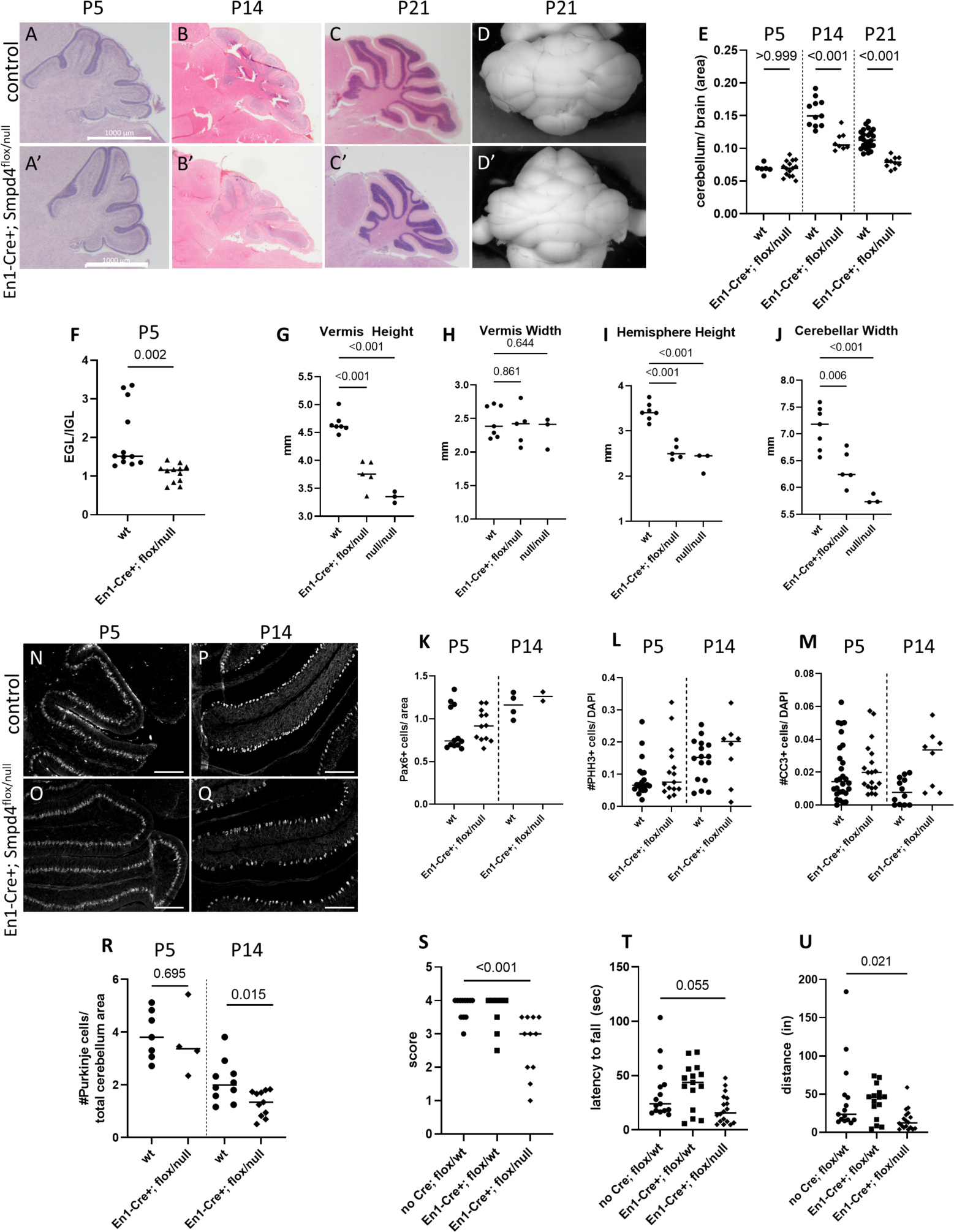
*Smpd4* is required for proper Purkinje cell support of postnatal cerebellar development. *En1*-Cre mediated *Smpd4* deletion leads to a progressively underdeveloped cerebellum (A-D’, quantified in E) and shows premature exit of granule precursor cells from the external to internal granule layer at P5 (F). By P21, the height of the vermis and hemispheres, and the overall width of the cerebellum, is drastically reduced although vermian width appears unaffected (G-J). The number of granule precursor cells in the external granule layer marked by PAX6 (K) and cell division rates (pHH3, L) are not different at P5 and P14. Cell death (CC3, M) is increased at P14 but not P5. A specific loss of Purkinje cells is seen between P5 and P14 (N-Q, quantified in R). The deletion mice have corresponding cerebellar behavioral phenotypes including hindlimb clasping (S) and increased latency to fall and decreased distance traveled on a rotarod (T,U).

Next, we assessed which cell type(s) are perturbed leading to the smaller cerebellar size and quantified the number of cells in both the EGL and internal granule layer (IGL). *En1-Cre+; Smpd4^flox/null^* animals have a lower ratio of EGL to IGL cells at P5, suggesting that more granule precursor cells are already exiting the proliferative EGL and populating the nascent IGL (**Fig 3F**). We hypothesized this is due to premature differentiation and/or migration of the granule precursor cells. We quantified these PAX6-positive cells and found no difference in their number between control and conditional knock-out animals at P5 or P14 (**Fig 3K**). Similarly, pHH3-positive proliferative cells are not reduced (**Fig 3L**). We did find an increase in apoptotic cells (CC3-positive) at P14, but not P5 (**Fig 3M**). We noted that most of the dying cells were in the Purkinje cell layer. While Calbindin-positive Purkinje cells are present in normal numbers at P5, they fail to survive and are markedly decreased at P14 (**Fig 3N-Q, quantified in R**).

Purkinje cell loss often causes ataxic behaviors. We scored P21 *En1-Cre+; Smpd4^flox/null^* mice for behavior indicating cerebellar ataxia (Lukacs et al., 2020) and found they exhibit hindlimb clasping (**Fig 3S**). We also tested them at six weeks of age on a rotarod to further quantify cerebellar ataxia (Deacon, 2013). As expected, *En1-Cre+; Smpd4^flox/null^* mice performed worse than their littermates, with shorter latency to fall and shorter total distance traveled on the rod (**Fig 3T,U**). No sex bias was observed with either behavioral test. Overall, this data supports a model wherein *Smpd4* is required for postnatal survival, but not initial generation, of Purkinje cells. Purkinje cell loss results in a smaller cerebellum and ataxic behavior in the mice.

### Forebrain-specific deletion of Smpd4 does not cause microcephaly

*Emx1* is expressed in the forebrain lineage as early as E10.5 (Gorski et al., 2002). We hypothesized such a specific deletion may reveal a microcephaly phenotype given the patient phenotypes, while circumventing lethality. *Emx1-Cre+; Smpd4^flox/null^* mice survive in Mendelian ratios to weaning but are not microcephalic (**Supp Table 6, Supp Fig 3A-J**). To test this ablation may not have been early enough in forebrain development, we next utilized *Foxg1*-*Cre* which is expressed in the telencephalic lineage beginning at E8.75 (Hébert and McConnell, 2000; Kawaguchi et al., 2016). *Foxg1-Cre+; Smpd4^flox/null^* mice likewise survive to weaning (**Supp Table 7**) but also do not exhibit microcephaly (**Supp Fig 3K-R**). Because human *SMPD4* patients have microcephaly, we hypothesized there is either a compensatory mechanism in mouse, or that human neurodevelopmental differences explain this disparity.

### Neutral sphingomyelinase genes do not compensate for each other in brain development

Given our finding that conditional *Smpd4* loss in the mouse forebrain does not lead to microcephaly as in humans, we hypothesized that neutral sphingomyelinase genes could act redundantly in the mouse. *Smpd3* and *Smpd4* are the two family members with highest expression in the developing brain (Fietz *et al*., 2012; Krut *et al*., 2006). We further detailed *Smpd3* mRNA expression across forebrain and cerebellar development (E12.5-P21). *Smpd3* is expressed at low levels in the developing cortex by E12.5 (data not shown), but unlike *Smpd4,* is restricted to upper cortical layers throughout forebrain development at E14.5 and E18.5 (**Supp Fig 4A,B**). It is also expressed in the cerebellar primordium (**Supp Fig 4C,D**). We observed no upregulation in mRNA expression of either neutral sphingomyelinase in the absence of the other (**Supp Fig 4E-P**). SMPD3 protein was previously shown to localize to the Golgi apparatus (Stoffel et al., 2005), the endoplasmic reticulum (ER), and the cell membrane (Piët et al., 2022). We did not observe localization of SMPD3 to the GM130-positive Golgi apparatus (**Supp Fig 4Q**). SMPD3 is also localized to the Calnexin-positive ER (**Supp Fig 4,R**) but not to the Lamin-positive nuclear membrane (**Supp Fig 4S**). Our RNA expression and protein subcellular localization data show some, but not complete overlap, of SMPD3 and SMPD4. We therefore genetically tested the hypothesis that there may be some functional redundancy between *Smpd3* and *Smpd4* during CNS development.

Previous studies showed that *Smpd3* deletion in mouse models causes juvenile dwarfism, delayed puberty, skeletal growth inhibition due to disruption of the Golgi secretory pathway of chondrocytes, and progressive cognitive impairment similar to Alzheimer’s disease (Stoffel *et al*., 2016; Stoffel et al., 2019; Stoffel *et al*., 2005). This allele of *Smpd3* was made on a C57BL/6×129Sv background and analysis was reported as a congenic C57BL/6 (B6) line. The independently derived *fragilitus ossium Smpd3* allele recovered from a forward genetic screen has osteogenesis imperfecta (‘brittle bone’ disease), shortened and bent long bones, tooth development issues, and partially penetrant postnatal lethality. We were unable to obtain either of these alleles and therefore generated a novel *Smpd3* deletion allele on a B6 background (**Supp Fig 5A-B**, hereafter referred to as *Smpd3^del^*). We found that deletion of *Smpd3* causes perinatal lethality, as we never recovered any live *Smpd3^del/del^* mice at birth (P0) (**Supp Fig 5C, Supp Table 8**). E18.5 embryos have severe skeletal abnormalities (**Supp Fig 5D-G**) including shortened mandibles (**Supp Fig 5H**) and long bones (radius, humerus, and scapula, **Supp Fig 5I-K**). However, we did not note any skeletal abnormalities in E18.5 *Smpd4^null/null^* embryos (**Supp Fig 6**). *Smpd3^del/del^* embryos do not exhibit any obvious cortical abnormalities at E14.5 or E16.5 (**Supp Fig 5L-N**). On a CD1 background, *Smpd3^del/del^* mice likewise exhibit complete perinatal lethality (**Supp Table 9**) and similar severe skeletal abnormalities (**Supp Fig 5O-V**). Given the earlier lethality in this model compared to previous studies, we performed whole-genome sequencing of an *Smpd3^del/del^* homozygote to ensure the fidelity of our allele. We found the genomic deletion was indeed specific to *Smpd3* and there were no other appreciable deletions on chromosome 8 (**Supp Fig 7**). Our data show that *Smpd3* is required for postnatal survival and skeletal development but is, on its own, dispensable for brain development.

Accordingly, *Smpd3^del/del^; Smpd4^null/null^* double knockout (dKO) mice do not survive birth and we noted some lethality at late embryonic stages (n= 8 of 19.75 expected at E16.5-E18.5, p=0.001, **Supp Table 10**). However, we did not observe any embryonic structural cortical abnormalities in dKO animals (**Supp Fig 4S-T**) and found no difference in cortical thickness (**Supp Fig 4U-V**). Overall, we saw no evidence for functional compensation in mouse brain development between these two neutral sphingomyelinase genes.

### Human patient iPSC models exhibit cell death and decreased proliferation

Mouse neurogenesis is a model for many aspects of human neurogenesis, but the human forebrain is markedly more complex. Therefore, we hypothesized that a human model might reveal specific consequences for cortical neurogenesis upon loss of *SMPD4*. We acquired fibroblasts from an *SMPD4* patient and reprogrammed them into iPSCs. We also created an *SMPD4* knock-out iPSC line (*SMPD4* KO) using CRISPR/Cas9 (**Supp Fig 8**). Neural rosettes are two-dimensional structures of neural progenitor cells (NPCs) with radial organization similar to the *in vivo* cortex (Fedorova et al., 2019; Knight et al., 2018; Zhang et al., 2001). We generated neural rosettes from these iPSCs and found several abnormalities in both patient and KO rosettes. Brightfield images after eight days in culture show that *SMPD4* patient and KO rosettes exhibit disrupted morphology with cells missing from the center (**Fig 4A**). Specifically, they have a smaller area and diameter (**Fig 4E,F**), fewer PAX6-positive neural progenitors (**Fig 4B**), a fifty percent reduction in proliferation levels (pHH3, p<0.05, **Fig 4C,G**), and a two-fold increase in CC3-positive, apoptotic cells (p<0.001, **Fig 4D,H**).

**Figure 4:**
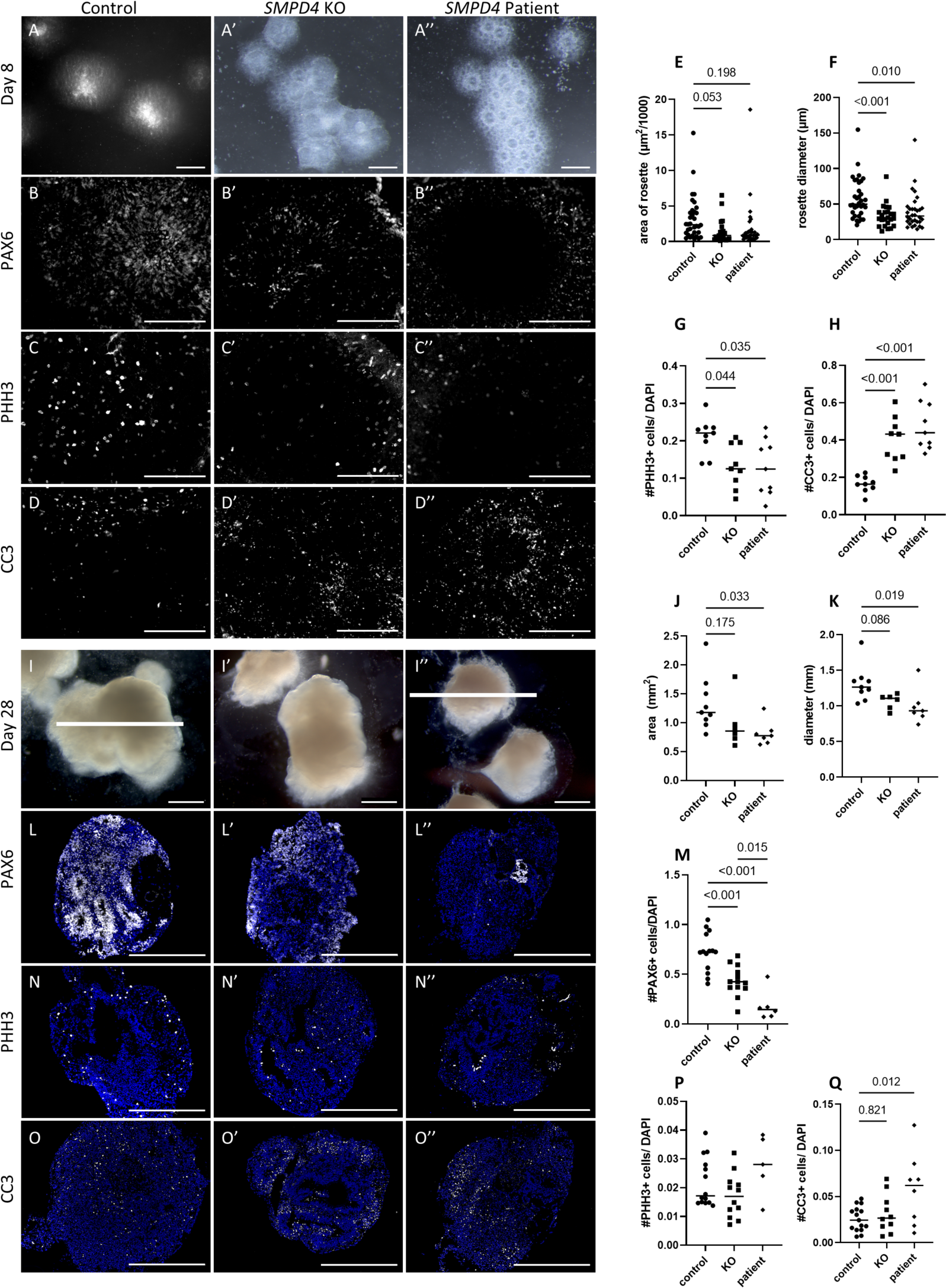
Human iPSC models show reduced neural progenitor cells, decreased proliferation, and increased cell death. Neural rosettes generated from control iPSCs at eight days in culture show normal rosette formation (A), whereas *SMPD4* patient and knockout rosettes are structurally abnormal (A’-A’’) and smaller (E,F). Immunohistochemistry for neural progenitors (PAX6), cell proliferation (pHH3), and cell death (CC3) (B-D and quantified in G,H) indicating a decrease in proliferative cells and increase in apoptotic cells in both *SMPD4* patient and knockout rosettes. Neural organoids generated from iPSCs likewise show that KO and patient organoids are smaller than control (I-K). KO and patient organoids have a distinct loss of PAX6-positive progenitor cells (L,M). The number of pHH3-positive dividing cells is unchanged (N,P), but apoptosis is increased in patient organoids (O,Q). Scale bars in A,I,L,N,O indicate 500 μm. Scale bars in B-D indicate 200μm. White bar in I and I’ are the same size to highlight smaller size of patient organoids.

While neural rosettes are a good two-dimensional model for early neurogenesis, neural organoids permit three-dimensional study of human brain development at later stages *in vitro* (Lancaster et al., 2013). After 28 days *in vitro*, neural organoids generated from *SMPD4* patient and KO iPSCs were smaller than control organoids (p=0.019 and p=0.086, respectively, **Fig 4I**, quantified in **J,K**). Interestingly, we detected an even more severe loss of PAX6-positive NPCs in the organoid model as compared to the neural rosettes, with patient organoids containing virtually no NPCs (**Fig 4L**, quantified in **M,** p<0.001). Proliferation levels were similar (**Fig 4N, P**), and cell death is less severe than in the neural rosettes although still increased in patient-derived organoids (**Fig 4O,Q,** p=0.012). We hypothesize that the proliferation and cell death differences between these two iPSC models is because they model different stages of cortical development. An early loss of NPCs as seen in the neural rosette model disrupts early neurogenesis, explaining a more severe size difference persisting in the organoid model of later stage development.

Premature loss of NPCs, decreased proliferation, and increased cell death are common molecular mechanisms of human microcephaly (Insolera et al., 2014; Jayaraman *et al*., 2016; Kaindl et al., 2010; Kumar *et al*., 2009; McIntyre *et al*., 2012). Because fewer NPCs are present, proliferate less, and fail to survive, many fewer total neurons are produced. We propose a combination of these mechanisms explain the microcephaly seen in *SMPD4* patients. We hypothesize the difference between our mouse and human iPSC models indicates a species-specific use of *SMPD4* in cortical development.

### SMPD4 loss leads to cell-specific primary cilia defects

Ceramide depletion has previously been shown to disrupt primary ciliogenesis (Wang *et al*., 2009) and cilia are crucial for normal brain development (Lee and Gleeson, 2011; Park et al., 2019). We hypothesized that loss of SMPD4 would decrease ceramide availability at the primary cilium. The primary cilia in cultured mouse E14.5 forebrain neurons appear comparable to wild-type in number and length (**Fig 5A-D**). Because we have shown that the *En1-Cre+; Smpd4^flox/null^* mouse cerebellar defect is due to Purkinje cell death, we also looked specifically at these cells. Interestingly, while the number of cilia is unaffected, *En1-Cre+; Smpd4^flox/null^*Purkinje cell cilia are significantly longer at both postnatal stages we assayed (P5 and P14, **Fig 5E-H**).

**Figure 5:**
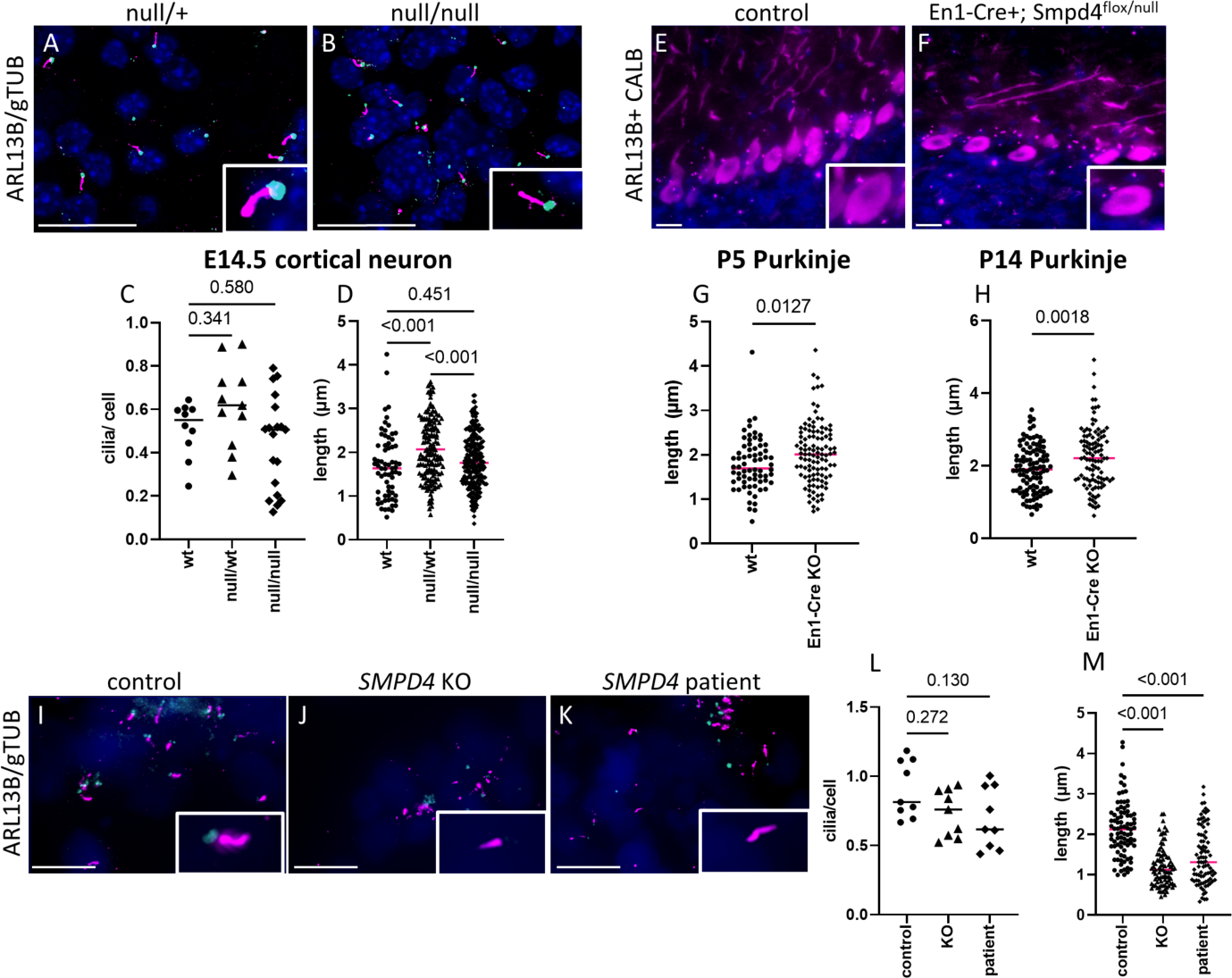
Primary cilia are disrupted by *SMPD4* loss. Primary cilia in neurons cultured from heterozygous *Smpd4^null/wt^* and homozygous *Smpd4^null/null^* E14.5 cortex are highlighted by ARL13B in the ciliary membrane and gamma tubulin at the basal body (A,B). Cilia appear to be unaffected in number or length (C,D). Tissue sections of postnatal mouse cerebellum stained with ARL13B and Calbindin (E,F) show that at P5 and P14, the number of primary cilia is unchanged in *En1-Cre+; Smpd4^flox/null^* animals (G,I) but cilia are longer (H,J). Cilia in human iPSC-derived neural rosettes (K-M) are present in normal numbers (N) but are significantly shorter (O) in *SMPD4* KO and patient. Scale bars in A,K-M= 10µm.

Cilia in human *SMPD4* KO and patient iPSC rosettes are found in normal numbers but are shortened and dysmorphic (control length=2.13 µm, KO= 1.19 µm, patient= 1.45 µm, p<0.001, **Fig I-M**). *SMPD4* KO and patient organoids also have shortened, dysmorphic primary cilia without a significant decrease in number (control length= 2.17 µm, KO=1.42 µm, patient= 1.31 µm, p<0.001, **Supp Fig 9**). The observed dysmorphisms were not uniform, but we noticed that some KO and patient rosette cilia have bulbous distal tips.

### Loss of Smpd4 does not lead to an accumulation of total sphingomyelin or decline in total ceramide in mouse brain

Neutral sphingomyelinases function in the recycling portion of the sphingolipid biosynthesis pathway to produce ceramide and phosphocholine from sphingomyelinase. The cell can use either this ‘recycling’ pathway or *de novo* synthesis but the majority primarily utilize the recycling pathway (Gillard et al., 1998). Thus, we hypothesized that loss of SMPD4 sphingomyelinase activity will cause a sphingomyelin buildup and/or ceramide decrease in the cell which could explain SMPD4 phenotypes. To test this, we performed mass spectrometry for sphingomyelin and ceramide species of *Smpd4* control and homozygous null E18.5 mouse brains in triplicate. Surprisingly, we found no overall change in sphingomyelin or ceramide in the cortex or cerebellum of mouse brain tissue (**Supp Fig 10**). These results are similar to those seen in *Smpd2* and *Smpd3* KO mice which also do not exhibit a change in sphingomyelin levels (Stoffel *et al*., 2005). Numerous studies have suggested that the activity of sphingomyelinases is highly specific to the organelle(s) in which they are localized. Therefore, perturbation of one sphingomyelinase may lead to organelle-specific changes in sphingolipid content rather than a tissue-or even cell-level change (Airola and Hannun, 2013; Hannun and Obeid, 2008; 2018).

### SMPD4 cilia defect is rescued by exogenous ceramide treatment

To test the effect of ceramide on promoting cilia growth, we treated human iPSC lines with exogenous C16 ceramide (**Fig 6A-M**). Treatment with GW4869 or FB1 results in fewer cilia in control iPSCs (**Fig 6N**), confirming that ceramide depletion causes a cilia defect. Interestingly, GW4869 has no effect on KO or patient cells, consistent with its known role as an nSMase inhibitor in cells that have already lost SMPD4 function (Luberto et al., 2002) (**Supp Fig 11).** *SMPD4* KO and patient cells do not have fewer cilia than control cells, and supplementation with ceramide did not significantly affect the number of primary cilia (p=0.668 and p=0.225, respectively, **Fig 6O**). However, ceramide supplementation dramatically increased cilia length in *SMPD4* KO and patient cells (1.61 µm to 2.19 µm for patient, 1.43 µm to 2.32 µm for KO, p<.001, **Fig 6P**). We therefore conclude that loss of functional SMPD4 leads to changes in local ceramide availability at the cilia, resulting in shortened and presumably dysfunctional cilia.

**Figure 6:**
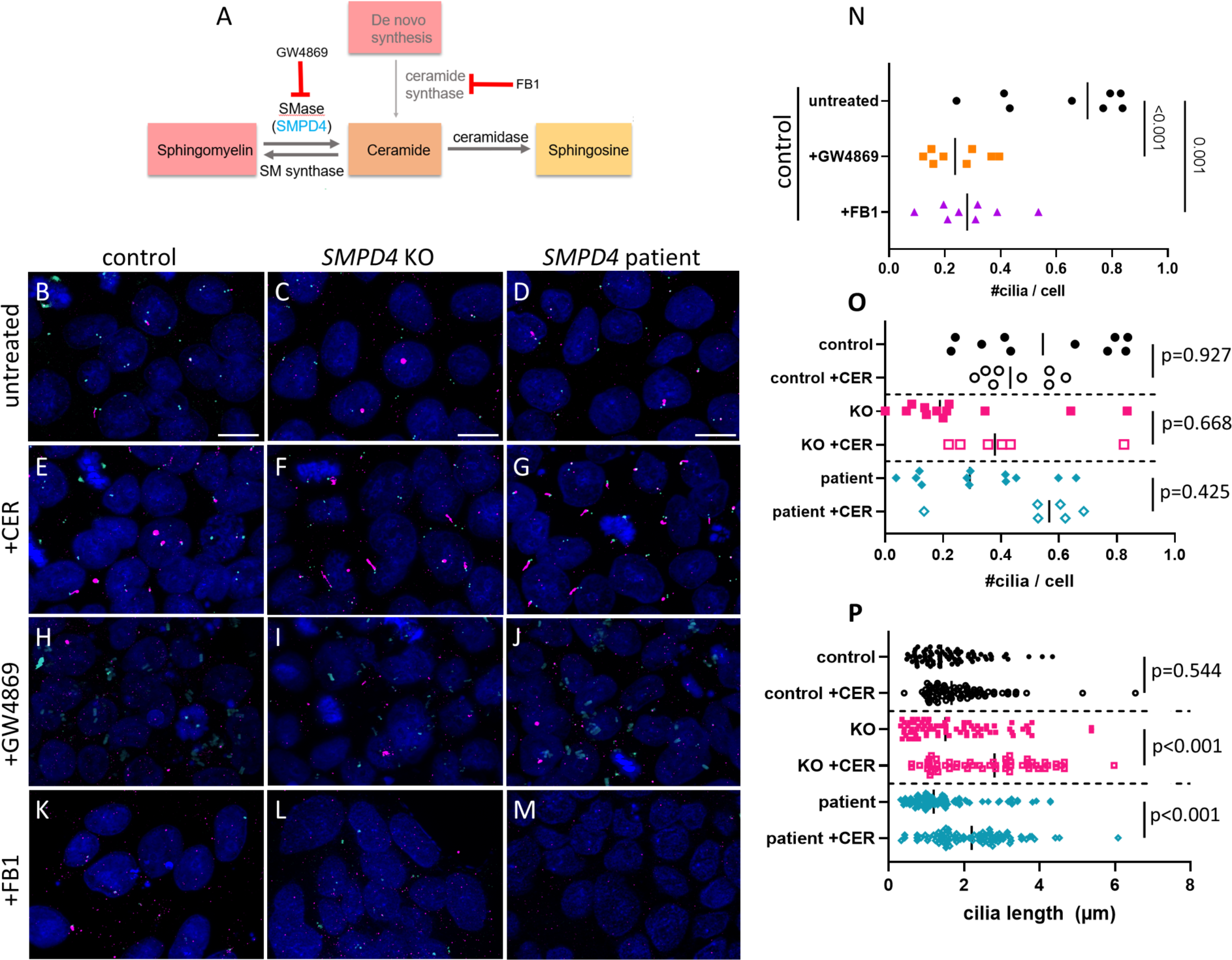
Cilia shortening in *SMPD4* models is rescued by *in vitro* ceramide treatment. iPSCs were treated with PBS, C16 ceramide, GW4869, or FB1 (A-M). Treatment with GW4869 or FB1 decreases number of cilia in control iPSCs (N). Treatment of *SMPD4* KO and patient iPSCs with ceramide does not impact the number but does increase the length of primary cilia per cell (O,P). Scale bars in B-D= 20µm.

### SMPD4 may impact molecular signaling pathways

Primary cilia are known to be crucial for proper signal transduction of several key developmental pathways including SHH and WNT. We hypothesized that loss of *SMPD4* and disrupted cilia function would have a transcriptional effect on developing neural tissues. To test this, we performed RNA sequencing of iPSCs and neural rosettes from control, *SMPD4* KO, and *SMPD4* patient conditions. Our analysis yielded 67 differentially expressed genes in iPSC KO and 2,222 genes in iPSC patient compared to iPSC control, and 74 genes in neural rosette KO and 2,463 genes in neural rosette patient compared to NR control (Fold change ≥ 1.5, FDR <10%, **Supp Fig 12A-D**). As the control line is isogenic to *SMPD4* KO but not *SMPD4* patient cells, we generally noted that there were not many differentially expressed genes common between KO and patient (**Supp Fig 12E**), potentially obscuring our further analysis.

Accordingly, standard pathway analysis failed to identify common functional themes among differently expressed genes. Given the changes in cilia, we specifically examined key genes in the SHH and WNT pathways. SHH signaling is largely unaffected (**Supp Fig 12F-H**). We did note several WNT target genes were highlighted by differential expression analysis and that WNT signaling is generally upregulated (**Supp Fig 12I-M**). WNT signaling is crucial for many distinct stages of neural development, suggesting an intriguing possible molecular consequence for loss of *SMPD4*.

## Discussion

We have demonstrated that *SMPD4* is critical for brain development in humans. We present a mouse model of loss of *Smpd4* which recapitulates human cerebellar hypoplasia but not microcephaly. Human iPSC models display cellular phenotypes consistent with disruption of forebrain development, including a novel ciliary dysfunction. Shortened primary cilia in *SMPD4* models are rescued by addition of exogenous ceramide and show evidence for dysregulated WNT signaling. Together, these results suggest that *SMPD4*-mediated sphingolipid metabolism regulates brain and primary cilia development.

### Human and mouse model discrepancies

We did not find a molecular mechanism leading to the reduced forebrain observed in the *SMPD4* germline null mouse model, although these mice rarely survive postnatally. Additionally, forebrain specific deletion of *SMPD4* did not lead to microcephaly. Finally, a zebrafish *smpd4* loss-of-function model showed no developmental or neural abnormalities (Smits *et al*., 2023). However, we found strong evidence for disrupted forebrain development with our *in vitro* human iPSC models. Overall, these results strongly suggest a species-specific difference in the role of *SMPD4* in forebrain development. The human cortex is markedly expanded relative to other species. Developing human brains undergo several additional rounds of neural progenitor proliferation chiefly driven by radial glia-like progenitors in the outer subventricular zone (Hansen et al., 2010), a cortical region found in primates but not mouse. Further, the mouse brain is naturally lissencephalic (smooth) whereas the human cortex undergoes cortical folding in order to accommodate the increased number of neurons in gyrencephalic species (Subramanian et al., 2020). If the outer subventricular zone neural progenitors are particularly perturbed by loss of *SMPD4*, we would expect to see a more severe cortical phenotype in human than in mouse.

### Purkinje cells in cerebellar development

Purkinje cells are the only output cell of the cerebellum and form a circuit with the cerebellar cortex via their axonal extensions to the deep nuclei. In mouse, they are produced from E10.5-E13.5 and migrate until E17.5, when they form a monolayer underlying the EGL (Altman and Bayer, 1985a; b; Rubenstein and Rakic, 2013). The interaction between Purkinje cells and granule cell precursors (GCPs) is essential for granule cell proliferation and foliation (Corrales *et al*., 2004; Wechsler-Reya and Scott, 1999). Postnatally, Purkinje cell survival and dendritic development significantly contribute to the cerebellar cytoarchitecture. Purkinje cells are particularly susceptible to cell degeneration and death and their loss leads to several neurodegenerative disorders including autosomal dominant cerebellar ataxias (Huang and Verbeek, 2019). Loss of *Acer3*, an alkaline ceramidase which breaks down ceramide and sphingosine-1-phosphate, causes Purkinje cell degeneration in adult mice via accumulation of long-chain ceramides and complex sphingolipids (Wang et al., 2015). Mice with a deletion in ceramide synthase (*CerS1*) have smaller Purkinje cells with underdeveloped dendritic arbors and progressively lose Purkinje cells starting at P21 (Zhao et al., 2011). The *Smpd4* null mice exhibit cerebellar hypoplasia consistent with postnatal loss of Purkinje cells. Purkinje cells in the null mouse are born in normal numbers and are present at the beginning of postnatal proliferation of GCPs but fail to survive, with apoptosis seen between P14-P21. Because GCPs are most highly proliferative from P3-P15, this suggests that loss of the Purkinje cells (and their dendritic arbors) results in reduced trophic support for GCPs during this period and a smaller cerebellum. Why loss of *SMPD4* causes Purkinje cells to fail is a fascinating question for future work.

### Smpd3/4 have the same biochemical function but are not redundant

We hypothesized that the lack of microcephaly in the *Smpd4* forebrain conditional knockout mice could be due to *Smpd3* compensatory sphingomyelinase activity in the mouse forebrain. However, this was not supported by *Smpd3/Smpd4* double knockout experiments, nor did *Smpd3* and *Smpd4* expression in the mouse forebrain completely overlap.

No human variants in *SMPD3* have yet been published. No homozygotes with predicted loss of function variants, and very few homozygotes for any missense variants, are present in the Genome Aggregation Database (gnomAD), suggesting *SMPD3* variants are not tolerated well in humans. Reported phenotypes for *Smpd3* mouse models differ. The *Smpd3^fro^* fragilitas ossium mouse displays postnatal lethality on a C57BL/6 background, runting, shorter long bones due to endochondral ossification abnormalities and decreased mineralization of cortical bones, and dentinogenesis imperfecta (Khavandgar et al., 2011). An independent *Smpd3* deletion mouse shows low embryonic lethality with postnatal growth retardation, short stature, deformed long bones with delayed ossification, and Alzheimer’s-like neurological phenotypes in adulthood (Stoffel *et al*., 2005; Stoffel et al., 2018). The *Smpd3* deletion mouse allele reported here exhibited complete perinatal lethality with shortened and malformed long bones but no evident cortical brain abnormalities. Interestingly, previous work noted that unlike SMPD4, SMPD3 does not associate with nucleoporins (Piët *et al*., 2022). This is confirmed by our subcellular localization data which shows that SMPD4 is present on the nuclear membrane, but SMPD3 is not. Therefore, the forebrain development difference between the human and mouse models is not explained by redundant neutral sphingomyelinase activity in the mouse. Collectively, these studies suggest unique subcellular roles for *SMPD3* and *SMPD4*.

### The role of SMPD4 and ceramide in centrosome dynamics and the cell cycle

During mitosis, the centrosomes separate by moving along the nuclear envelope to orient the mitotic spindle (Tanenbaum and Medema, 2010). Both the nuclear envelope and nuclear pore complexes break down and reassemble dynamically during mitosis (Antonin et al., 2008), which is mediated by nucleoporins (Hawryluk-Gara et al.; Kutay et al., 2021). We and others have documented the subcellular localization of SMPD4 to the ER, nuclear envelope, and the pericentriolar region during mitosis (Smits *et al*., 2023). Therefore, we hypothesize that the localization of SMPD4 and/or ceramide to the nuclear envelope is essential for interacting with the centrosome. Cell cycle disturbances have previously been shown in *SMPD4* patient fibroblasts as well as HEK293T and HeLa si*SMPD4* cells (Atilla-Gokcumen et al., 2014; Magini *et al*., 2019; Smits *et al*., 2023). SMPD4 associates with nucleoporins including NUP35 and NDC1 (Piët *et al*., 2022). Delays in nuclear pore complex insertion and envelope closure were seen with SMPD4 knockdown (Smits *et al*., 2023). Nuclear pore formation defects and variants in nuclear pore proteins are known to cause microcephaly (Carapito et al., 2019; Fujita et al., 2018; Ravindran et al., 2021; Ravindran et al., 2023; Rosti et al., 2017; Sandestig et al., 2020). Nucleoporin proteins localize to the base of the primary cilium in addition to the nuclear envelope (Kee et al., 2012). This all suggests SMPD4 localization to the nuclear membrane and centrosome contributes to its molecular role.

The centrosome also serves as the basal body for the primary cilium. The cilium is primarily known as a sensory organelle crucial for the cell to respond to intercellular signaling but also affects cell cycle progression (Perry and Hannun, 1998). Additionally, because ceramide is required at the basal body for primary ciliogenesis and the cilium is resorbed during S phase, primary cilia shortening is caused by cell cycle defects (Seeley and Nachury, 2010). We propose a novel role for SMPD4 in controlling ceramide distribution to the nuclear envelope and primary cilium and have shown that it is important for ciliogenesis. Future work will determine the molecular mechanism by which SMPD4 impacts the cell and ciliogenesis cycles. A tantalizing possibility is that an effect on WNT signaling leads to corticogenesis deficits.

### Local ceramide distribution may be key to SMPD4 activity

Ceramide is the precursor of many complex sphingolipids (Sandhoff and Kolter, 2003). Ceramide is synthesized primarily via the ‘hydrolysis’ pathway (Gillard *et al*., 1998) in the cell membrane, *de novo* in the smooth ER, and rarely through the ‘salvage’ pathway in the lysosome. Although *SMPD4* knockdown or loss-of-function does not lead to accumulation of sphingomyelin or depletion of ceramide at the tissue level (Atilla-Gokcumen *et al*., 2014; Smits *et al*., 2023) (and this study), organelle localization of these sphingolipids is key to their activities (Airola and Hannun, 2013). Localized lipid distribution in organelle membranes, such as ceramide, changes the membrane curvature (Arya et al., 2022). This study also suggests that nSMase activity is needed for nuclear envelope buds to release and form cytosolic vesicles. Because ceramide biogenesis is thus specific to certain organelle membranes, we hypothesize that nuclear membrane loss of SMPD4 could lead to a local deficit in ceramide, perturbed local concentrations and compromised transport to the plasma membrane. This would also result in a deficit of ceramide available at the plasma membrane where acetylated tubulin associates with ceramide-rich platforms for primary ciliogenesis (Tripathi *et al*., 2021). Bulk ceramide supplementation is a candidate intervention but there is no established paradigm for this, and the above studies suggest that this may not alter local concentration in a therapeutic way.

### SMPD4 as a disease gene

In this work, we show a new role for *SMPD4* in the primary cilium. We link primary cilia length to ceramide, expanding our understanding of how deficits in neutral sphingomyelinase cause brain disorders. We have advanced the study of *SMPD4* as an interesting cause of a human structural brain disorder. This study suggests that modulating *SMPD4,* neutral sphingomyelinase function, and/or ceramide content in the cell would improve brain development in human patients.

## Acknowledgements

We are grateful to the Pluripotent Stem Cell Facility (Dr. Chris Mayhew) and Transgenic Animal and Genome Editing Core (Dr. Yueh-Chiang Hu) at Cincinnati Children’s Hospital and the Institute for Genomic Medicine Genomic Services Laboratory (Dr. Amy Wetzel and Ben Kelly) at Nationwide Children’s Hospital. We thank current and former members of the Stottmann lab, especially Bekah Rushforth, Elizabeth Bittermann, Lauren Blizzard, and Dr. Zakia Abdelhamed for their advice and support. This work was supported by the Cincinnati Children’s Hospital Medical Center for Mendelian Genomics and Therapeutics, Nationwide Children’s Hospital Abigail Wexner Research Institute, NIH R35GM131875 (R.W.S.), and NIH F31HD104350 (K.A.I).

## Author contributions

R.W.S. and K.A.I. conceived the study. K.A.I. and R.W.S. designed experiments. K.A.I. and B.C. performed experiments and analyzed the data. K.A.I. and R.W.S. wrote the original draft and edited the manuscript. All authors reviewed the manuscript before submission.

## Declaration of interest

The authors declare no competing interests.

## STAR Methods

### Generation of knockout mouse alleles

Mouse zygotes (C57BL6/N strain) were injected with 200 ng/µL CAS9 protein (Thermo Fisher, Waltham, MA) and 100 ng/µL *Smpd3*-specific sgRNA (TGGCCAGAGCAGGCTGCACG CGG, CAGGTCCTAAAGCAGCAGTC AGG, IDT), followed by surgical implantation into pseudo-pregnant female (CD-1 strain) mice (CCHMC TAGE Core, RRID:SCR_022642). The resulting live-born pups were weaned and used for additional PCR screening and mating.

*Smpd4* mice were obtained from the IMPC (RRID:SCR_006158, *Smpd4*^tm2a(KOMP)Wtsi^). The tm2a allele can be converted to tm2b (*null*) or tm2c (*flox*) in this system. We generated tm2b (*null*) mice by crossing tm2a with a germline Cre (EIIa-Cre, The Jackson Laboratory, RRID:SCR_004633), and tm2c *(flox*) mice by crossing with FLP carrier mice (The Jackson Laboratory). Conditional Cre alleles were from The Jackson Laboratory. *Smpd4^null/+^*; *Cre+* mice were crossed with *Smpd4^flox/flox^* to generate *Cre+/WT; Smpd4^flox/null^*; mutants, which we refer to as conditional KO mice. A list of mouse alleles used for this project is in **Supp Table 1.**

### Mouse husbandry

All animals were maintained through a protocol approved by Nationwide Children’s Hospital Medical Center IACUC committee (IACUC2021-AR1200067). Mice were housed with a 12-h light cycle with food and water *ad libitum*. Mouse euthanasia was performed in a carbon dioxide chamber followed by secondary cervical dislocation. Genotyping was performed via PCR and gel electrophoresis on a 2% agarose gel with the primers listed in **Supp Table 2**. Whole-brain and skeletal images were taken on a Zeiss Discovery V12 microscope (Zeiss, St. Louis, MO).

### Behavior Testing

Hindlimb clasping was measured on postnatal day (P) 21 by recording a 10-second video on a camcorder while the animal was suspended by the tail six inches from the cage floor. Two researchers blinded to genotype independently scored tail clasping on a 0-4 scale (Lukacs et al., 2019) and the average of the two measurements was used for statistical analysis. Motor coordination was tested at six weeks of age (n=9-10 animals per genotype) on a ROTOR-ROD™ (San Diego Instruments, San Diego, CA) with speed increasing from 0-50 rpm over a 2-minute testing period. The researcher was again blinded to genotype. The ROTOR-ROD™ system measured distance traveled (cm) and latency to fall (sec). An even number of males and females were tested per genotype for both behavior tests, and we note there was no statistical difference between sexes.

### Histology

Brains were dissected, fixed in formalin for 48h, washed in 70% ethanol, then dehydrated and paraffin-embedded by the morphology core. Blocks were sectioned on a microtome at 10µm (Sakura, Hayward, CA). Sections were placed on glass slides (Cardinal Health, Dublin, OH), baked >1 hour, and stained with hematoxylin and eosin using standard methods. Body and brain weights were obtained on a standard chemical scale. For cortical and cerebellar measurements, area in µm^2^ or length in µm was measured on ZEN 3.7 software. A minimum of 3 animals from at least 2 distinct litters were measured for each genotype.

### Skeletal preparation

Pups were collected at E18.5 and frozen. Skin and fat were removed from the embryos prior to fixation in 95% ethanol for 2-5d. The skeletons were stained with Alizarin Red and Alcian Blue and cleared with potassium hydroxide using standard methods (Behringer et al., 2014). Bone length measurements were taken with ZEN 3.7 software (n=3-4 animals per genotype).

### Immortalized cell culture

HEK293T cells (ATCC, Manassas, VA) were cultured in Dulbecco Modified Eagle Medium supplemented with 10% fetal bovine serum (FBS) and 1% penicillin/streptomycin at 37°C and 5% CO2. Cells were enzymatically detached from plates using 1 mL trypsin/EDTA (0.25%). Plasmid transfections used the Lipofectamine 3000 kit and manufacturer’s instructions (Thermo Fisher). A control GFP plasmid was used to assess transfection efficiency at 48 hours post-transfection (GFP-Nubp1, Okuno et al., 2010).

### iPSC culture

*SMPD4* patient fibroblasts were obtained from an affected individual with variant p.Glu124* (Magini *et al*., 2019) (Family 8) and reprogrammed to pluripotency by the CCHMC Pluripotent Stem Cell Facility (RRID: SCR_022634) using standard protocols. Control human iPSCs (Episomal hiPSC, Gibco) and patient-derived iPSCs were cultured in mTeSR media (STEMCELL, Cambridge MA) in Nunc plates (Fisher) on a matrix of Matrigel (Corning) dissolved in DMEM/F12. Cells were passaged every 7 days using Gentle Cell Dissociation Reagent (STEMCELL) and fed daily. For exogenous treatment experiments, iPSCs were plated onto Matrigel-coated coverslips in a 24-well plate. Cells were treated with Fumonisin B1 (Sigma Aldrich) at 30µM for 72 hours, GW4869 (Tocris Bioscience, Minneapolis MN) at 10µM for 72 hours, and/or C16 ceramide (Avanti Polar Lipids, Alabaster AL) at 2µM for 48 hours before being processed for immunocytochemistry.

The STEMdiff™ SMADi Neural Induction Kit (STEMCELL) was used to generate neural rosettes from high-quality iPSC colonies. Briefly, iPSCs were dissociated into single cells and plated into an Aggrewell 800 well (STEMCELL) at 10,000 cells per well, forming embryoid bodies. These were fed daily and replated on day five by filtering through a 37µM reversible strainer (STEMCELL) into a 24-well plate containing coverslips coated in Matrigel. On day 8, percent neural rosette formation was visually confirmed to be 75% or above before harvesting for immunocytochemistry.

Whole-brain neural organoids were generated from an adapted protocol based on Fair et al., 2023 (Fair et al., 2023). Cells were dissociated as above and resuspended in mTesR in 96-well ultra-low attachment plates on Day 0. On Day 10, embryoid bodies were transferred into 6-well plates and maintained until Day 28, when organoids were fixed overnight in 4% paraformaldehyde at 4°C and prepared for cryo-embedding and immunohistochemistry as described (Fair *et al*., 2023).

### CRISPR/Cas9 transfection

CRISPR guides were purchased from Synthego (Redwood City, CA). CRISPR editing was performed on control iPSCs using the Synthego CRISPR Lipofection protocol with GFP control as described above. Cells were serially diluted into a 96-well plate (10,000 cells/plate) for clonal isolation. Wells with only a single colony were grown to confluency, passaged into 6-well plates, and DNA was extracted for PCR to confirm CRISPR editing of the *SMPD4* locus. PCR product was purified with a DNA Clean & Concentrator kit (Zymo Research, Irvine, CA) and Sanger sequenced (**Supp Fig 6**). CRISPR guides and primers for genotyping are in **Supp Table 2.**

### Immunocytochemistry

Cells were plated onto coverslips in a 24-well tissue culture plate. They were fixed in 4% paraformaldehyde (PFA) for 15 minutes. Coverslips were immersed in 0.1% Triton-X 100 for 5 minutes prior to blocking to permeabilize cell membranes. Coverslips were blocked in 4% NGS for 30 mins and incubated at 4°C overnight in primary antibodies. Secondary antibody staining for 1 hour was followed by DAPI incubation for 15 min. Coverslips were sealed to glass slides with ProLong Gold Antifade Mountant (Thermo Fisher). Images were acquired on a Zeiss Axio Imager.M2 with Apotome 3 and an Axiocam 305 monochrome camera. Antibodies and concentrations for immunocytochemistry are listed in **Supp Table 3**. CC3 positive, PHH3 positive, and PAX6 positive cells from embryonic mice, neural rosettes, and organoids were counted in the NIS-Elements Analysis program (Nikon, Melville NY) with three images quantified for each of n=3 replicates. Calbindin positive cells were quantified by the same method but with one image from each animal (n=3-9 per genotype).

For analysis of primary cilia, cells were serum-starved in Opti-MEM Reduced-Serum Medium (Thermo Fisher) for 24 hours before fixation in methanol. Z-stack images were taken on the same Zeiss Apotome at 63x magnification and then a maximum intensity projection was produced from the Z-stack. Cilia were quantified with a NIS-Elements Analysis program, with ∼1000 cells per replicate per condition. Cilia length was measured in µm with ZEN 3.7 using the maximum intensity projection images, 20-30 cilia per image with n=3 replicates per genotype. The transfection experiment and cell counting were repeated (n=3) for each condition.

### Immunohistochemistry

After dissection, brains were fixed for 1-2 days in PFA at 4°C. PFA was replaced with 30% Sucrose for 2 days before brains were embedded in Optimal Cutting Temperature solution (Sakura) and stored at −80°C. 10µm sections were obtained for mouse brain and human organoid samples on a Leica CM 1860 cryostat, placed on glass slides, and stored at −20°C. Slides selected for IHC were pre-warmed at 42°C for 10-15 minutes. Antibody retrieval was performed as previously described (Bittermann et al., 2019). Blocking and secondary antibody processing was as described above.

### RNA in situ hybridization

Wild-type mouse embryos maintained on a CD1 genetic background were dissected at ages E12.5, E14.5, E18.5, P0, and P28 and fixed in formalin for 16-24 h; brains were sub-dissected for ages E18.5-P27. Tissue was washed in PBS, then dehydrated and paraffin-embedded by the morphology core. Paraffin blocks were sectioned at 5 µm, placed on glass slides, and baked at 60°C for 1 hour. Target retrieval steps outlined in the ACDBio (Newark, CA) RNAScope protocol were followed based on recommendations for brain tissue, then slides were dried at room temperature overnight. Hybridization and amplification steps were performed using the HybEZ oven set at 40°C, RNAscope Multiplex Fluorescent Reagent Kit V2 (323100), TSA Cyanine 3 Fluorophores (NEL744001KT) at 1:750, and *Smpd4* probe custom designed by ACDBio.

### Mass spectrometry

Liquid chromatography-electrospray ionization/ mass spectrometry (LC-ESI-MS/MS) was performed. Crude lipid extracts were prepared from mouse brain tissue using a single-phase extraction system of ethyl acetate:isopropanol:water. Along with internal standards, the extract was passed through an HPLC system coupled to a mass spectrometer with electrospray ionization capability (ESI/MS) with precursor ion scan and multiple reaction monitoring enabled. Each decomposed sphingolipid was then quantified by comparing to internal standards.

### RNA/WGS Sequencing

Total RNA was extracted from iPSCs or neural rosettes using the Trizol (Tri Reagent, Sigma Aldrich) protocol with post-extraction cleanup. Samples were kept on ice until storage at −80°C. RNA concentration was quantified via Nanodrop. DNA was extracted from tail clips with a DNEasy Blood & Tissue Kit (Qiagen, Germantown, MD) for whole genome sequencing. DNA concentration was quantified via Qubit. RNA and WGS sequencing were performed by the NCH Genomics Services Laboratory using standard Illumina protocols.

### Statistical analysis

Data plots and subsequent analyses were performed with Prism 9 (GraphPad, San Diego, CA). A student’s t-test was performed for experiments with two groups. An ANOVA with Tukey’s multiple comparison tests was performed for experiments with more than two comparisons. The p-values for these experiments are shown in the relevant figure. We report the statistical test values directly rather than assigning a significance symbol to provide all the data for the reader.

## Materials Availability Statement

All unique/stable reagents generated in this study are available from the lead contact with a completed materials transfer agreement.

## Supplemental Material

**Supplemental Figure 1:**
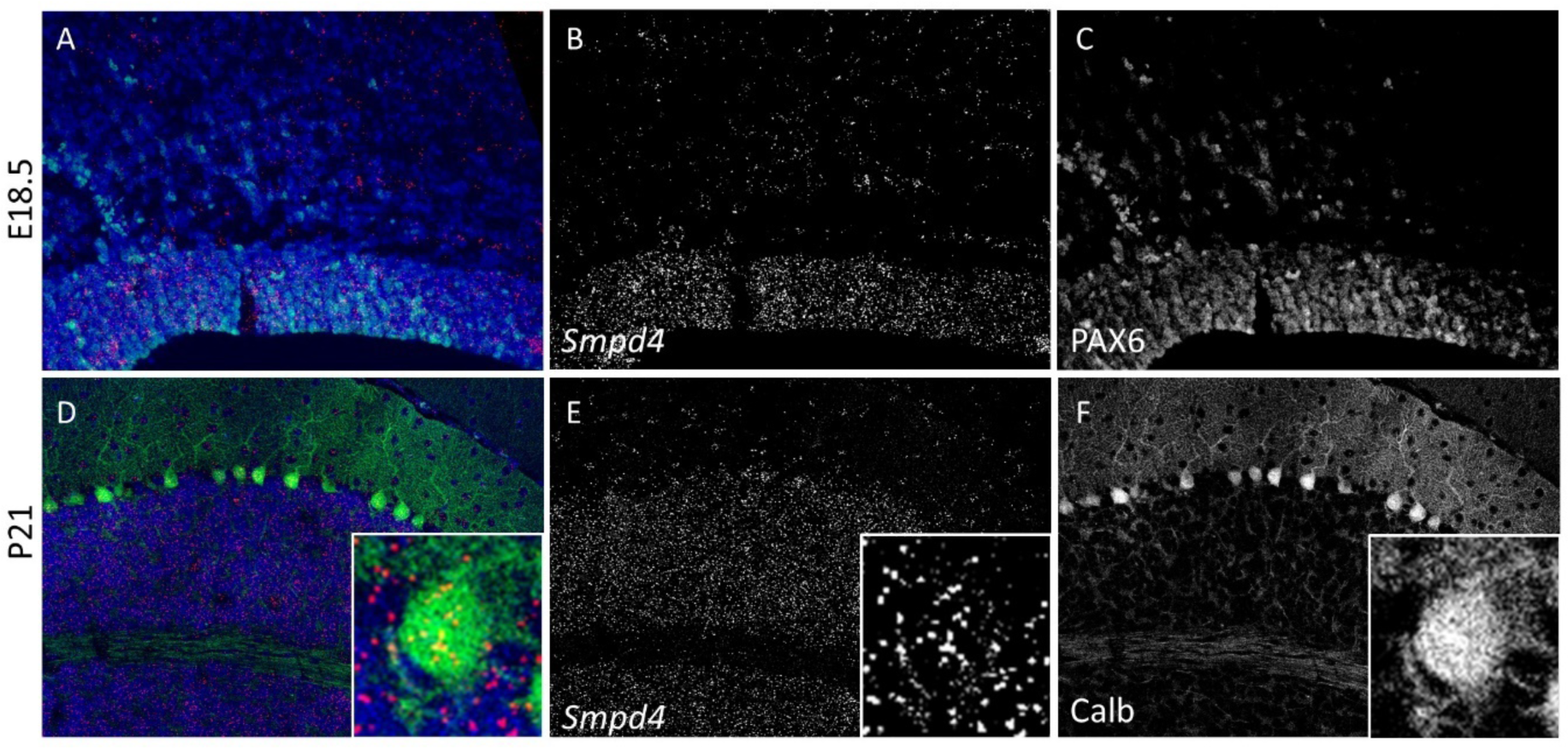
*Smpd4* mRNA colocalizes with cell type-specific markers. All images are *Smpd4* RNA *in situ* hybridization on one serial section overlaid with an immunohistochemistry antibody from the adjacent serial section, as in Figure 1D-F, J-L. At E18.5, *Smpd4*/ PAX6 colocalization is maintained and *Smpd4* becomes more restricted to the ventricular zone where PAX6-positive cells remain (A-C). In the cerebellum, *Smpd4* cerebellar expression remains in both the internal granule cell layer and in Purkinje cells marked by Calbindin to P21 (D-F).

**Supplemental Figure 2:**
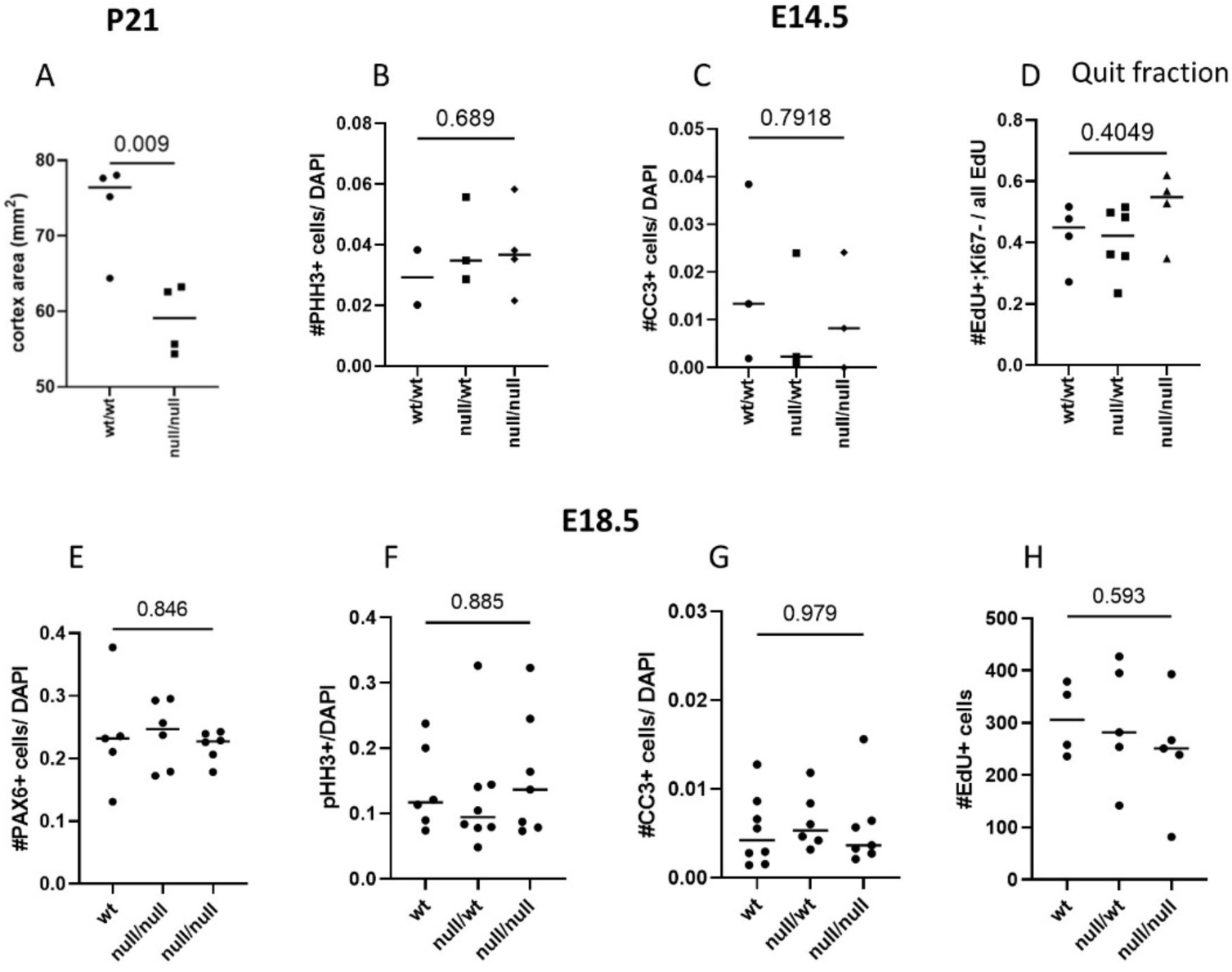
Normal proliferation and cell death in the *Smpd4* homozygous null forebrain. The cortical area of weaning stage *Smpd4^null/null^* animals was found to be smaller than their littermates (A) in addition to having a smaller body weight and failure to thrive. Next, forebrains from E14.5 *Smpd4* wild-type, heterozygous, and homozygous null animals were analyzed for several molecular markers. Quantification of pHH3-positive cells indicates no difference in proliferation (B). Likewise, the number of CC3-positive apoptotic cells in the ventricular zone (C), and EdU-positive divided cells labeled at E13.5 in cortex (D) are also unchanged. Similarly at E18.5, the number of PAX6-positive neural progenitors in the cortex (E) is unaffected, as are the number of pHH3-positive (F), CC3-positive (G), and E13.5 labeled EdU-positive cells (H).

**Supplemental Figure 3:**
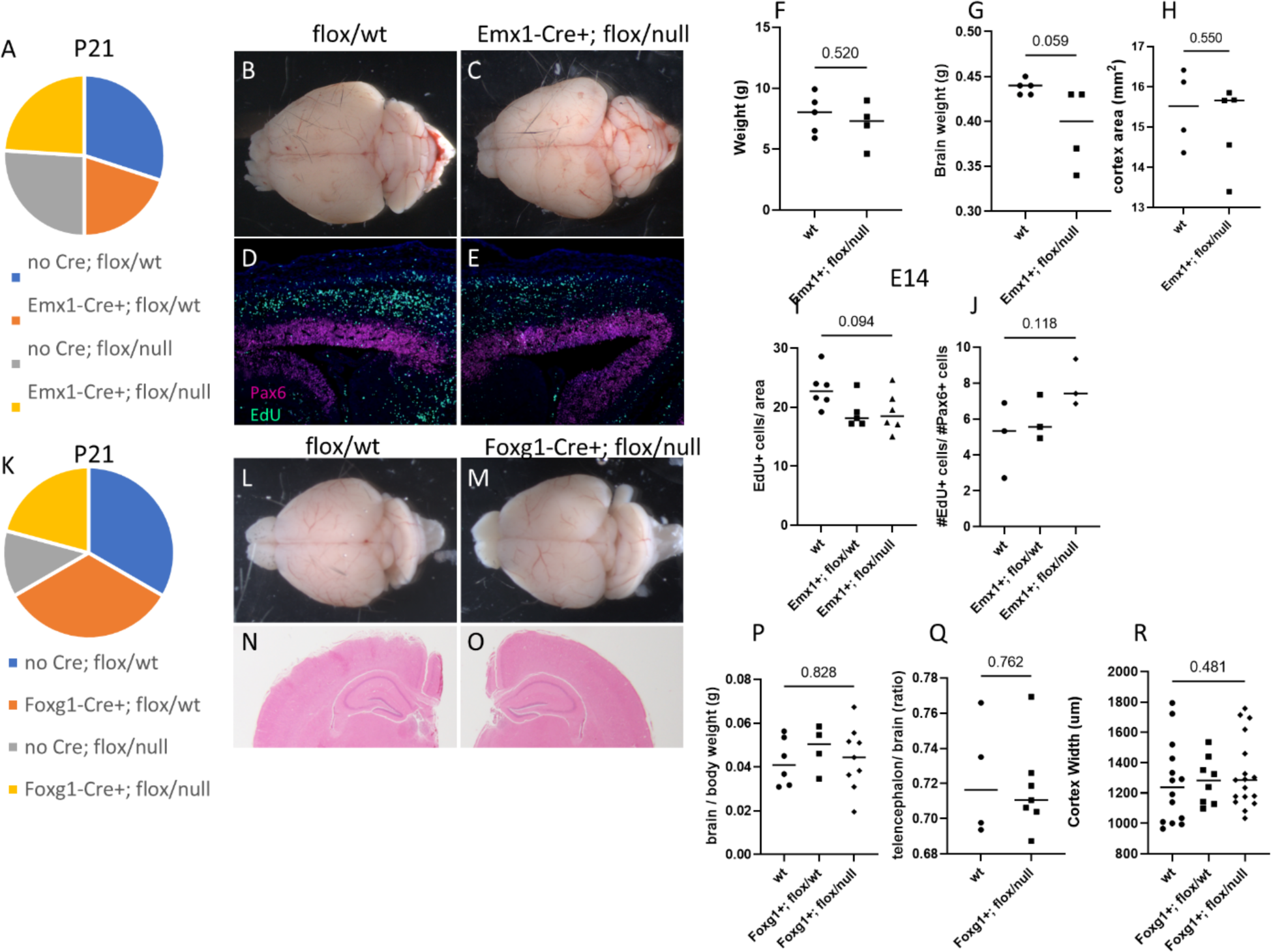
Conditional forebrain ablation of *Smpd4* rescues survival but does not recapitulate microcephaly. Emx1-Cre+; *Smpd4*^flox/null^ animals survive in Mendelian ratios at weaning (A) and do not exhibit any gross cortical abnormalities (B-C). They are the same size as their littermates (F) and demonstrate a reduction in brain weight (G), but not cortical area (H). Injection of pregnant females with EdU at E13.5 and dissection of embryos at E14.5 suggest normal cortical migration (D-E, quantified in I-J). Similar results were obtained for Foxg1-Cre+; *Smpd4*^flox/null^ mice (K-R).

**Supplemental Figure 4:**
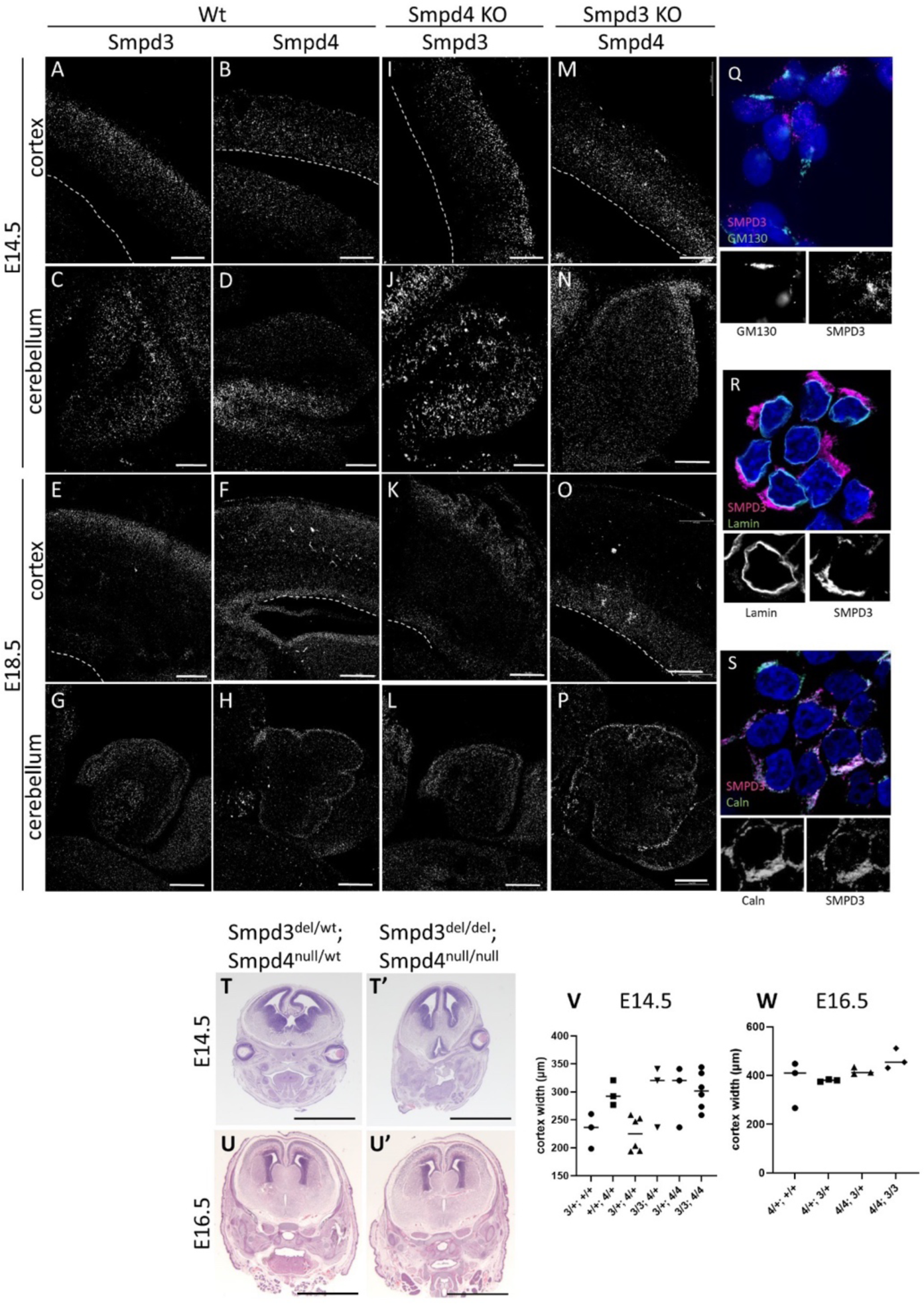
Neutral sphingomyelinases do not compensate for each other in mouse. RNA *in situ* hybridization shows that at E14.5 and E18.5, *Smpd3* is restricted to upper cortical layers in the forebrain (A,B) but is expressed throughout the cerebellar primordium (C,D). Compensation of neutral sphingomyelinases was tested by assaying *Smpd3* mRNA in the absence of *Smpd4* (*Smpd4^null/null^*, I-L) and *Smpd4* mRNA in *Smpd3^del/del^* cortex, finding no upregulation in mRNA expression in either case. Subcellularly, SMPD3 protein does not localize to the GM130-positive Golgi apparatus (Q) or Lamin-positive nuclear membrane (R) but is localized to the Calnexin-positive ER (S). Double knockout *Smpd3^del/del^; Smpd4^null/null^* animals do not exhibit any obvious craniofacial abnormalities (T,U), and their cortical width is unchanged at E14.5 and E16.5 (V,W).

**Supplemental Figure 5:**
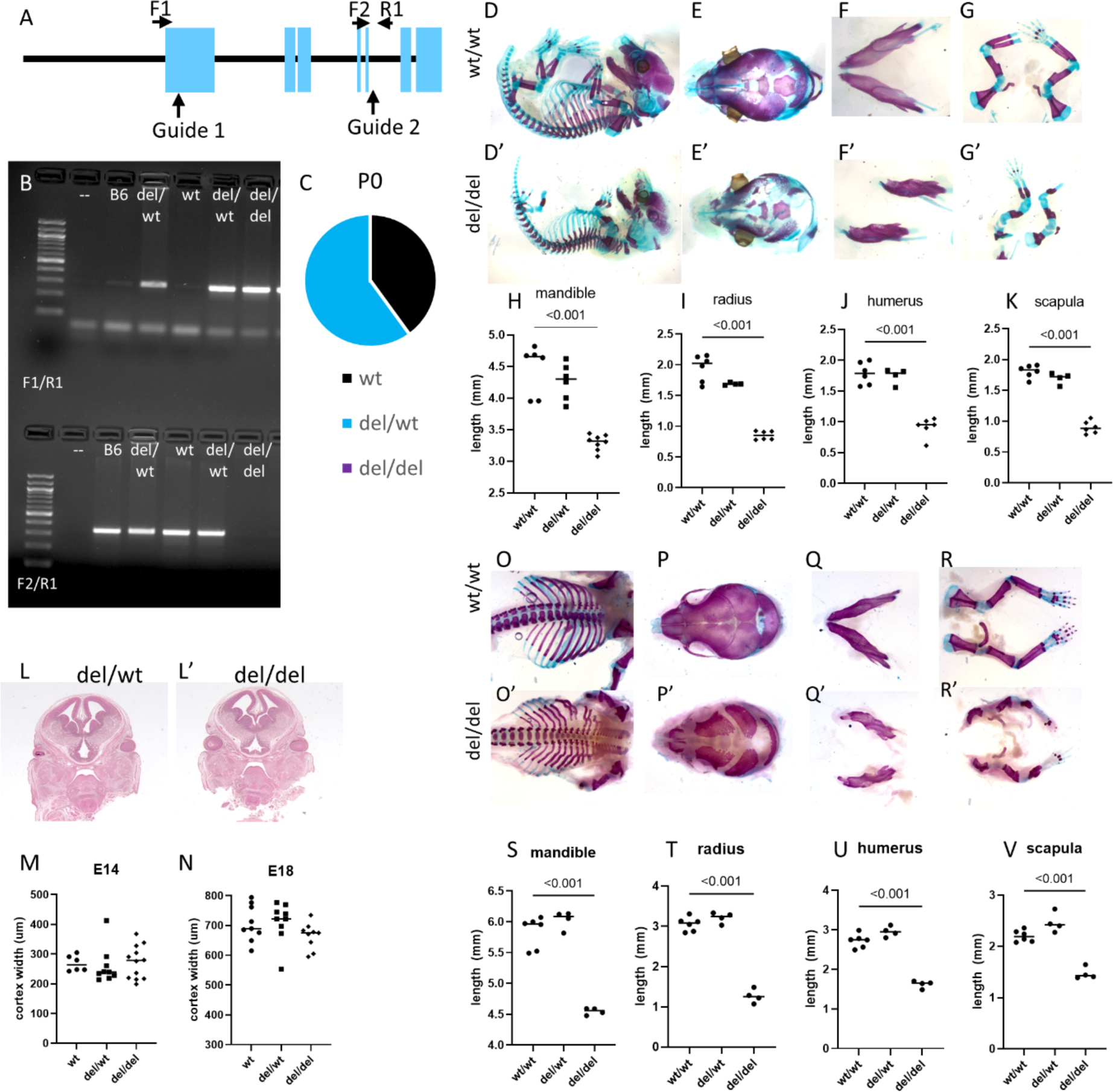
Generation of a novel *Smpd3* allele reveals requirement for postnatal survival and skeletal development but not brain development. CRISPR/ Cas9 guides were designed to induce a large deletion in *Smpd3* (A). Primers to genotype resulting animals are schematized (A) and results are shown in B. A PCR product of primers F1 and R1 indicates deletion of exons 1-5. *Smpd3* homozygous deletion animals do not survive at birth (C). Skeletal preps from E18.5 animals show overall dysmorphic and shortened bones in homozygous deletion animals (D-G, quantified in H-K). However, coronal sections at E14.5 (L-M) and E18.5 (N) do not show brain abnormalities or cortical thinning. We confirmed the phenotype is the same on an outbred CD1 strain (O-V).

**Supplemental Figure 6:**
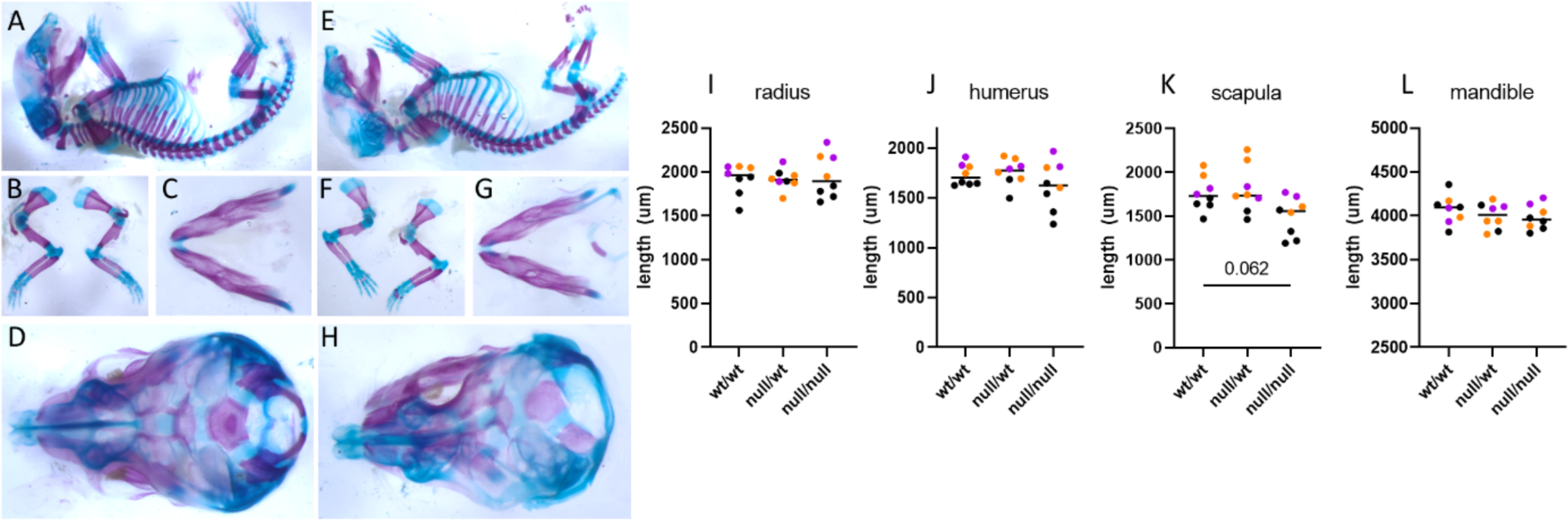
*Smpd4* mice do not exhibit skeletal abnormalities. E18.5 skeletons of *Smpd4* wild-type (A-D) and homozygous null (E-H) show no differences in bone (Alizarin Red) or cartilage (Alcian blue) appearance. The lengths of various long bones and the mandible are quantified in I-L, respectively.

**Supplemental Figure 7:**
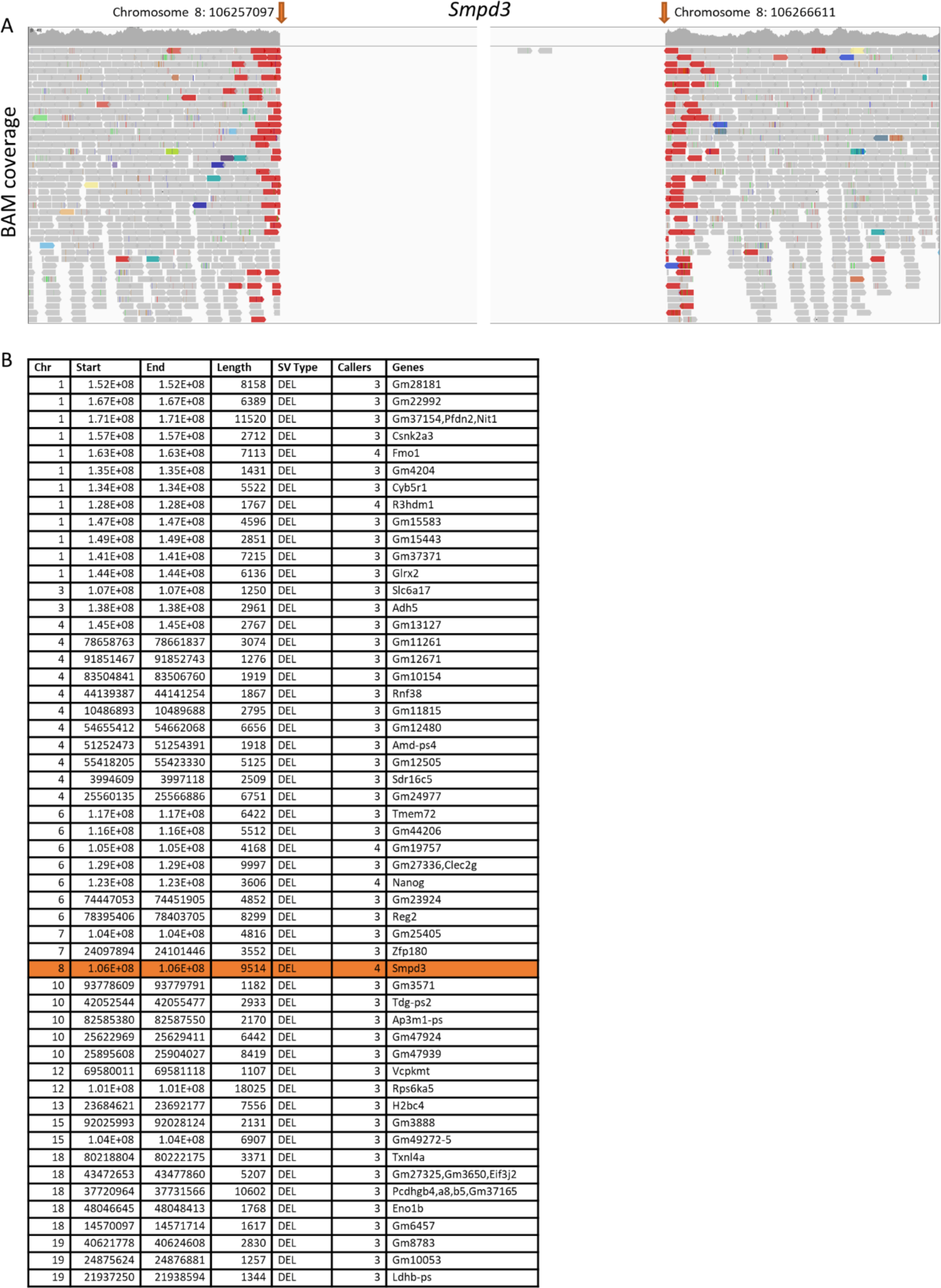
*Smpd3* homozygous mouse sequencing shows a specific large deletion. Concurrent with our Sanger sequencing confirmation of CRISPR/Cas9-mediated editing, whole genome sequencing of a *Smpd3^del/del^* mouse indicated a large 9514 base-pair deletion corresponding to exons 3-7 of *Smpd3*. Read sequence alignment is shown in A. There were no other appreciable large deletions/ structural variants on chromosome 8 (B).

**Supplemental Figure 8:**
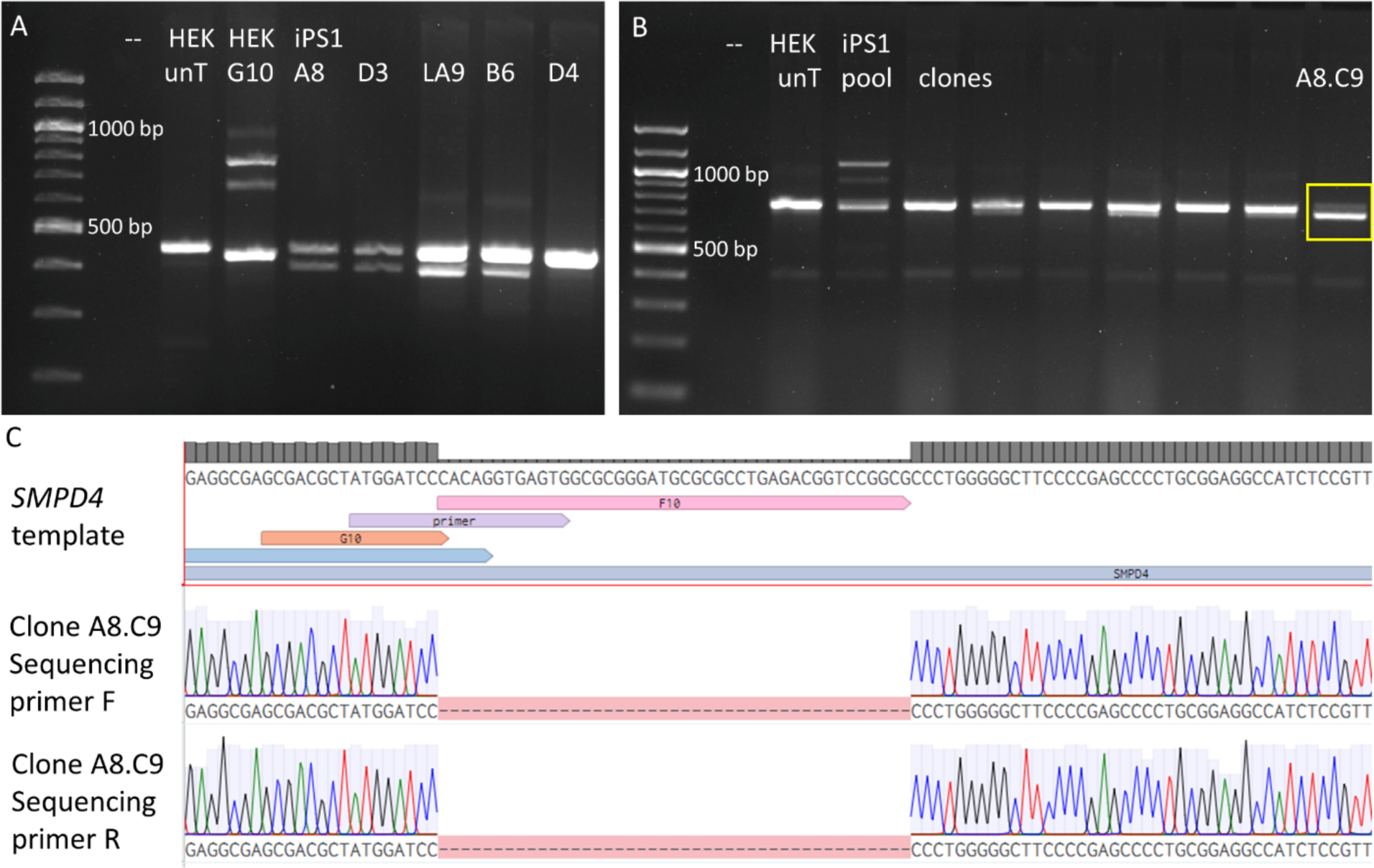
CRISPR Editing of *SMPD4* Locus in human iPSCs. CRISPR/Cas9 editing was used to disrupt the end of Exon 1 of *SMPD4*. CRISPR guides in an RNP complex designed by Synthego were transfected into control iPSCs. PCR of single clones showed a promising edited lower band for several clones including A8 (A) whereas clone D4 was unedited (HEK=HEK293T cell line; iPS1= control iPS line; unt= untransfected). Sanger sequencing for clone A8 showed a heterozygous 43 base pair deletion in *SMPD4*. The CRISPR experiment was repeated on this clone to edit the other allele. Sanger sequencing was performed with forward and reverse primers and each sequencing result was aligned to the GRCh38 human genome (see SMPD4 sequencing primer F and R, Supp Table 2). Alignment in Benchling confirmed a homozygous 43 base pair deletion in clone A8.C9 (C), which was used as our SMPD4 knockout (KO) line in the iPSC experiments in this manuscript.

**Supplemental Figure 9:**
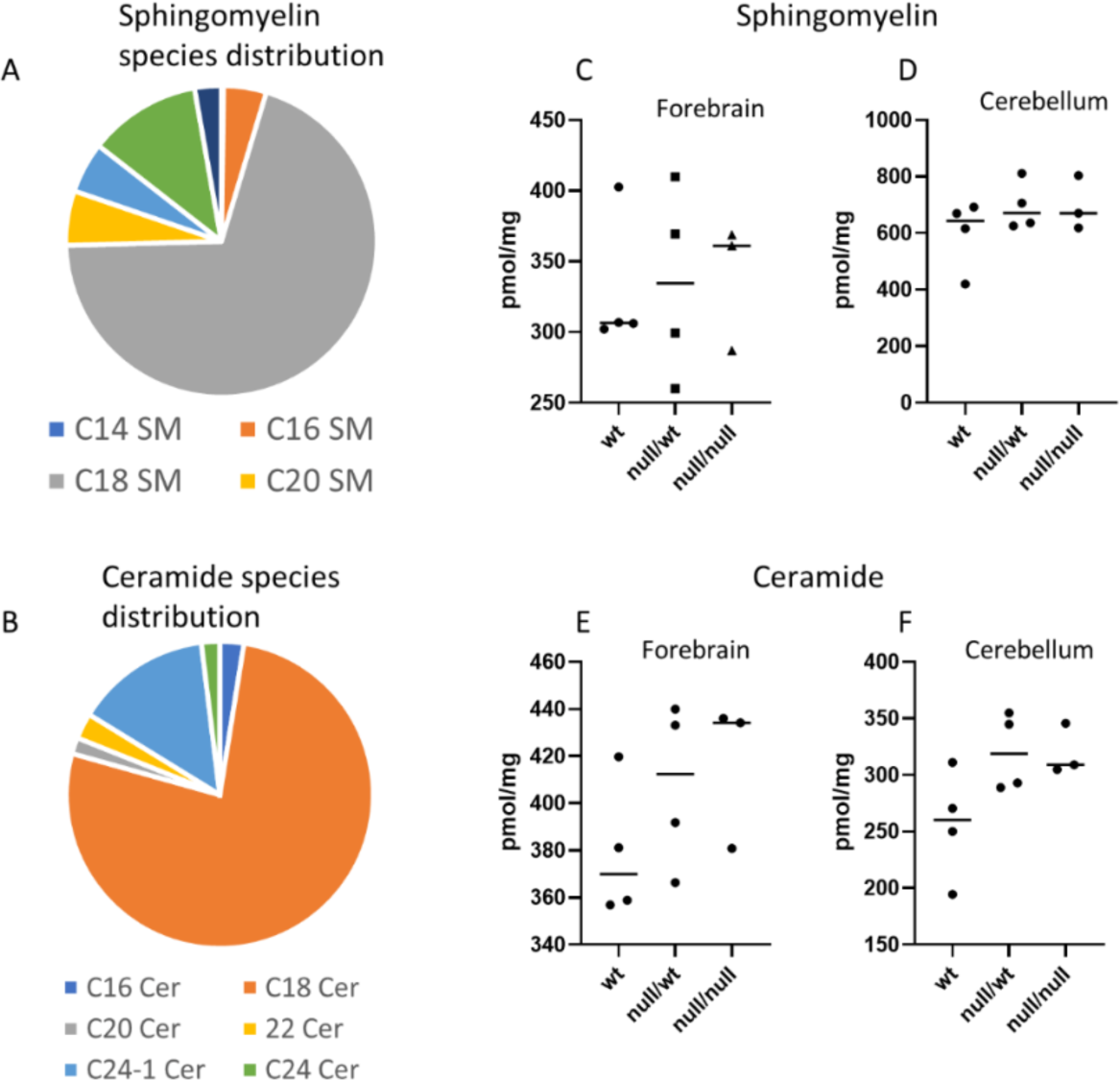
Primary cilia length in *SMPD4* models is also shortened in human iPSC-derived neural organoids. We generated three-dimensional neural organoids which model a later stage of human neural development than two-dimensional neural rosettes. At 28 days in vitro, control organoids exhibit widespread primary cilia lining each mini-ventricle (ARL13B in pink, A). *SMPD4* KO (B) and patient (C) iPSCs form neural organoids with abnormal cilia. Specifically, they do not have a significant decrease in the number of cilia per cell (n=2 images x 3 replicates, D), but are shortened in length (n=30 cilia x 3 replicates, E).

**Supplemental Figure 10:**
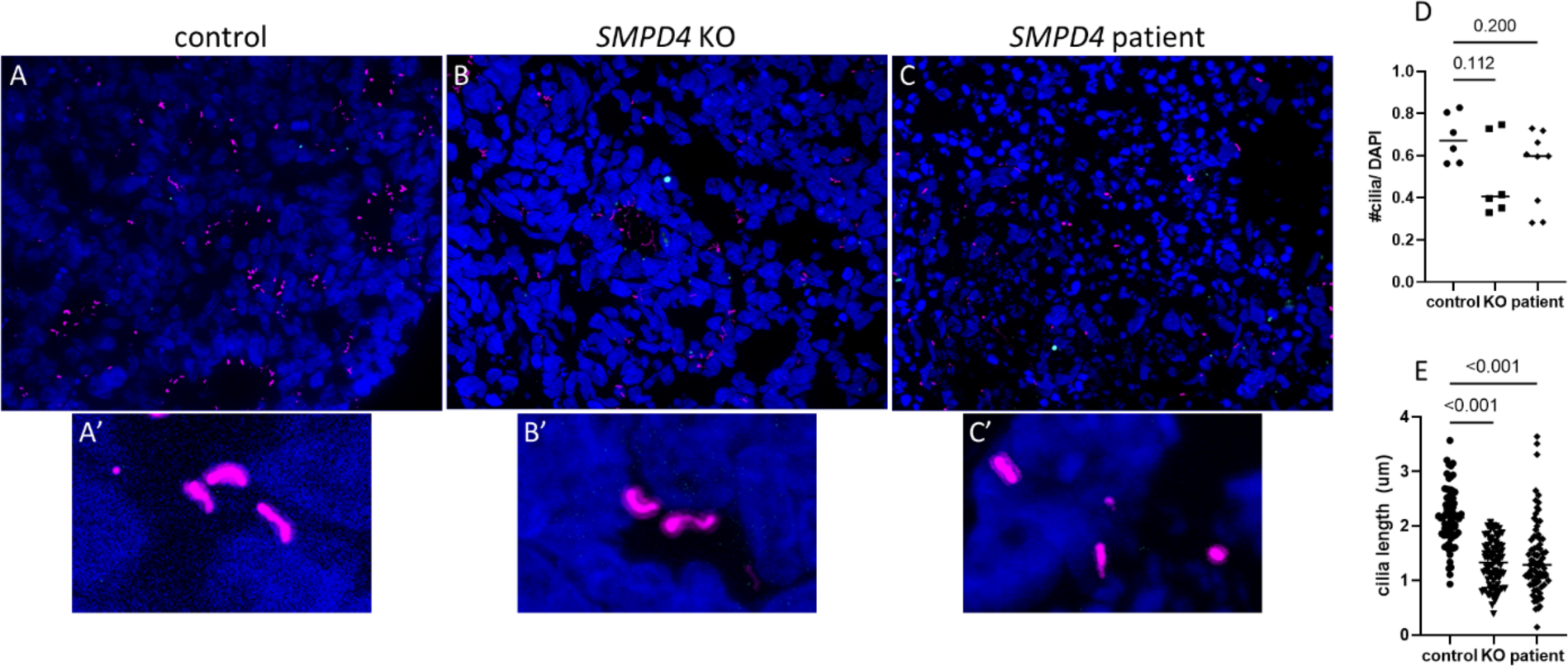
Sphingomyelin and ceramide levels are unchanged in *Smpd4* null mouse brain. Relative amounts of sphingomyelin (A) and ceramide (B) species in control E18.5 embryos. Mass spectrometry of E18.5 mouse brain tissue for sphingomyelin and ceramide results are summarized in C-D, E-F respectively. No differences are seen in overall amount of either sphingolipid in forebrain (C, E) or cerebellum (D, F).

**Supplemental Figure 11:**
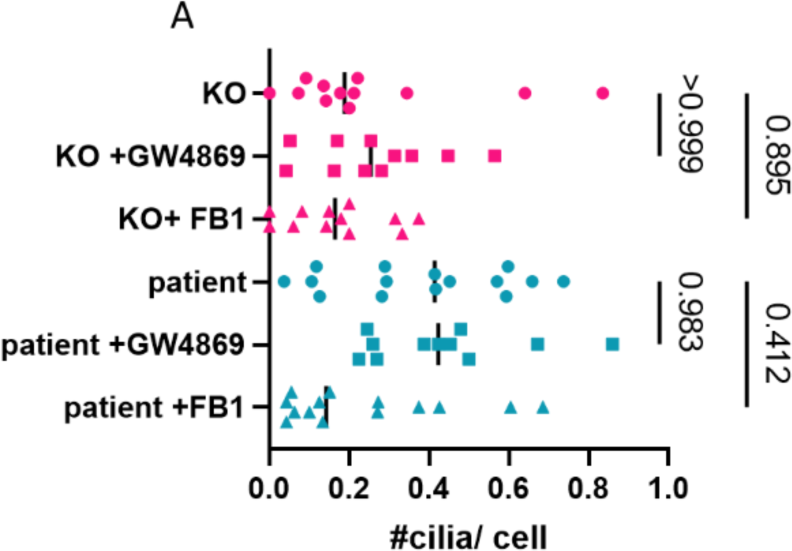
*SMPD4* KO and patient iPSCs are unaffected by blocking neutral sphingomyelinase activity. *SMPD4* KO and patient lines do not have decreased cilia number with GW4869 or FB1 treatment.

**Supplemental Figure 12:**
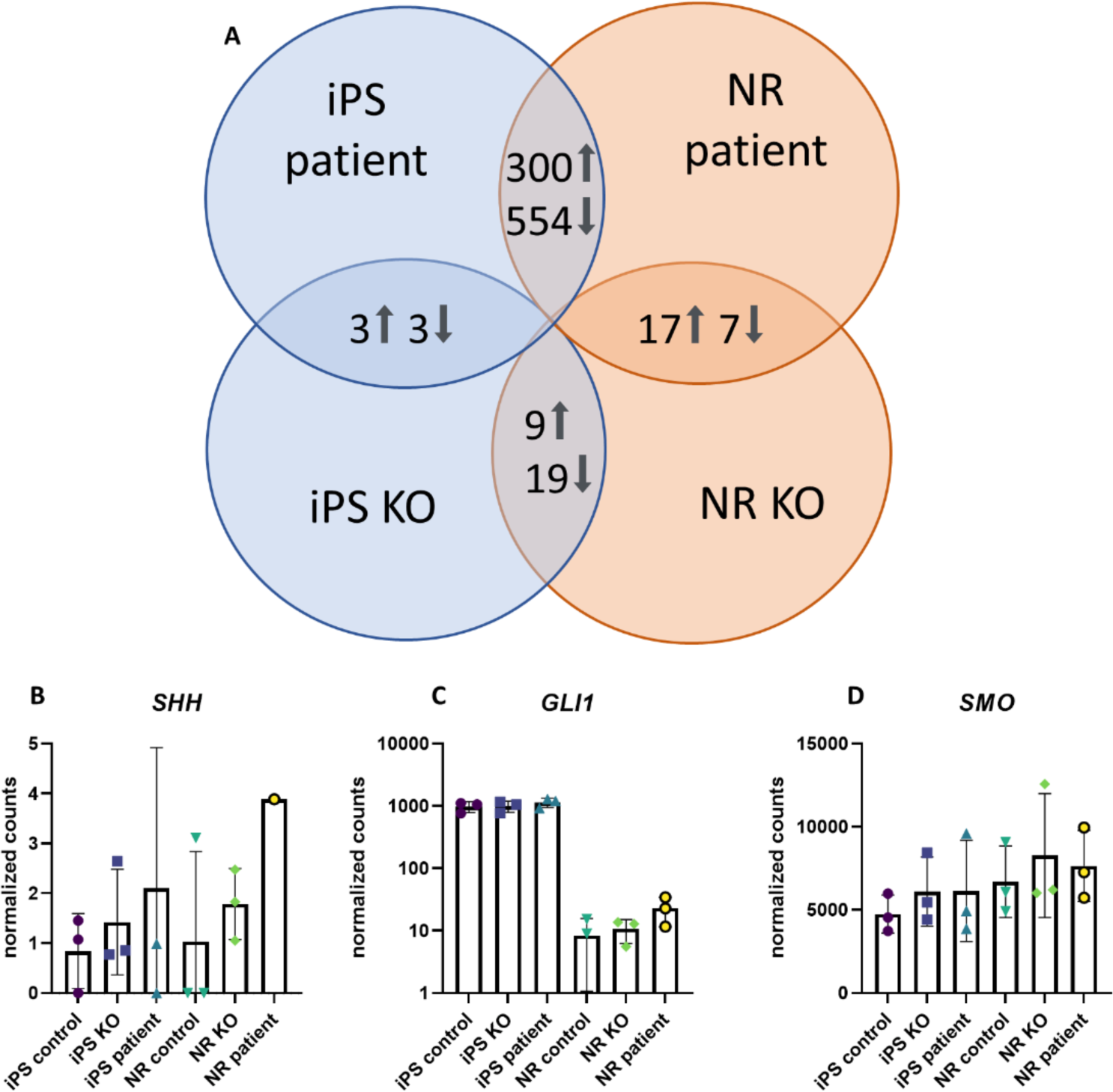
Loss of *SMPD4* does not impact SHH signaling, but may deregulate the WNT pathway. Our RNA sequencing differential expression analysis showed that there were not many differentially expressed genes common between KO and patient datasets (A). We took a deeper look at SHH pathway genes and found SHH signaling appears unaffected by loss of *SMPD4.* The *SHH* ligand itself is expressed at low levels across all samples and is upregulated ∼2 fold in patient neural rosette replicates only (B). *GLI1* and *SMO*, which are direct transcriptional targets of *SHH,* are unaffected (C,D). We also noted in our manual analysis that WNT signaling is upregulated. The central WNT pathway regulators *WNT3A* and *AXIN2* greatly increased in KO and patient NRs relative to control (I,J). We also identified upregulation of *APC2, WNT4,* and *WNT5A* (K-M).

**Supplementary Table 1:** Mouse Lines.

| Mouse Line | Source | RRID | Abbreviation |
| --- | --- | --- | --- |
| *Smpd4*<sup>tm2a(KOMP)Wtsi</sup> | IMPC | MMRRC_062656-UCD | *Smpd4*<sup>del</sup> |
| *Smpd4*<sup>tm2b(KOMP)Wtsi</sup> | n/a | n/a | *Smpd4*<sup>null</sup> |
| *Smpd4*<sup>tm2c(KOMP)Wtsi</sup> | n/a | n/a | *Smpd4*<sup>flox</sup> |
| B6.FVB-Tg(Ella-cre)<sup>C5379Lmgd/J</sup> | JAX #003724 | IMSR_JAX:003724 | Ella-Cre |
| FLP | JAX # 012930 | IMSR_JAX:012930 | FLP |
| B6.129S2-Emx1<sup>tm1(cre)Krj/J</sup> | JAX #005628 | IMSR_JAX:005628 | Emx1-Cre |
| En1<sup>tm2(cre)Wrst/J</sup> | JAX #007916 | IMSR_JAX:007916 | En1-Cre |
| B6.129T(SJL)-Foxg1<sup>tm1.1(cre)Ddmo/J</sup> | JAX #029690 | IMSR_JAX:029690 | Foxg1-Cre |
| *Smpd3* deletion | CCHMC TAGE | n/a | *Smpd3*<sup>del</sup> |

**Supplementary Table 2:**
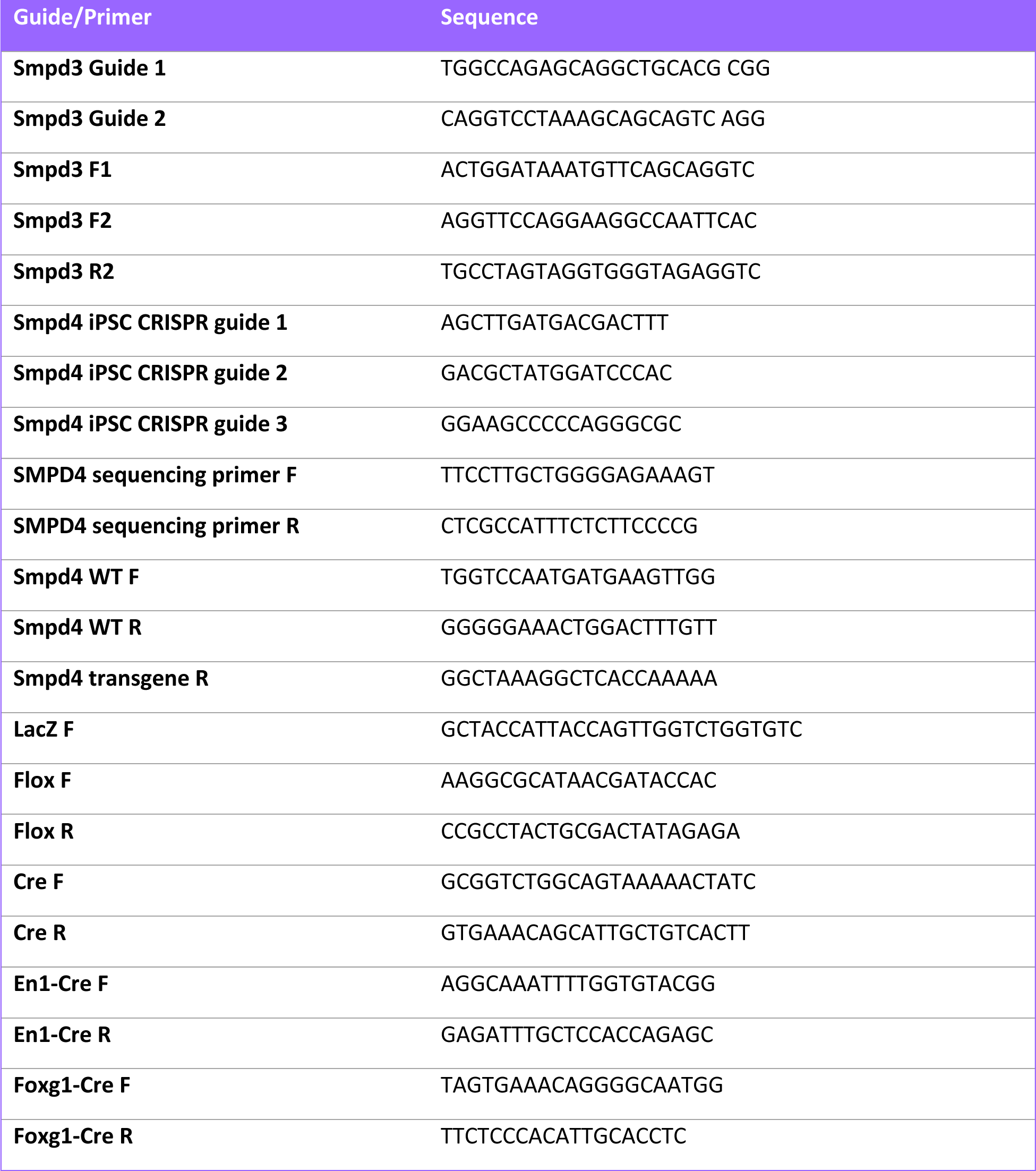
CRISPR Guide Sequences and Genotyping Primers.

**Supplementary Table 3:**
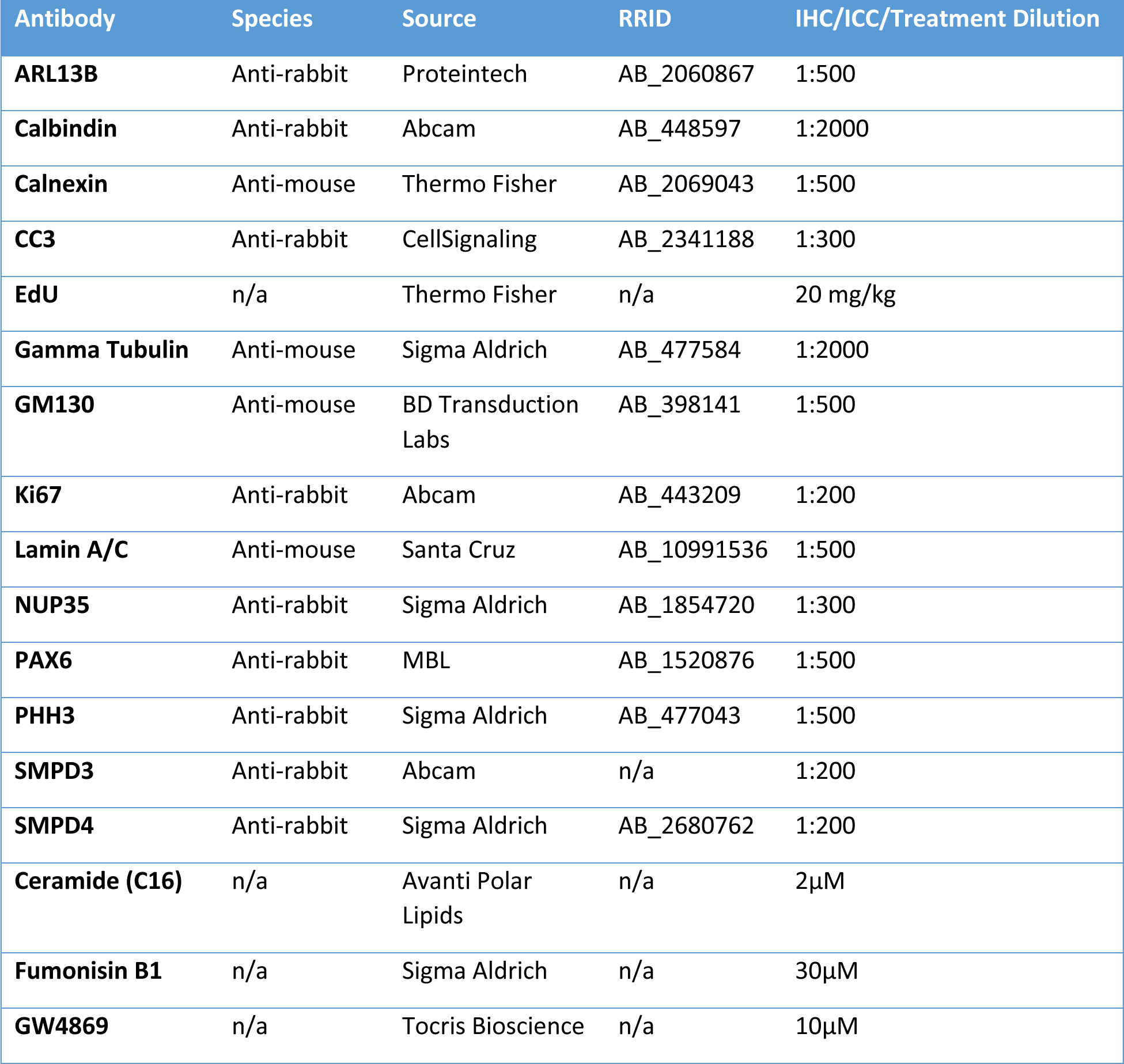
Antibodies and Compounds.

| Antibody | Species | Source | RRID | IHC/ICC/Treatment Dilution |
| --- | --- | --- | --- | --- |
| **ARL13B** | Anti-rabbit | Proteintech | AB_2060867 | 1:500 |
| **Calbindin** | Anti-rabbit | Abcam | AB_448597 | 1:2000 |
| **Calnexin** | Anti-mouse | Thermo Fisher | AB_2069043 | 1:500 |
| **CC3** | Anti-rabbit | CellSignaling | AB_2341188 | 1:300 |
| **EdU** | n/a | Thermo Fisher | n/a | 20 mg/kg |
| **Gamma Tubulin** | Anti-mouse | Sigma Aldrich | AB_477584 | 1:2000 |
| **GM130** | Anti-mouse | BD Transduction Labs | AB_398141 | 1:500 |
| **Ki67** | Anti-rabbit | Abcam | AB_443209 | 1:200 |
| **Lamin A/C** | Anti-mouse | Santa Cruz | AB_10991536 | 1:500 |
| **NUP35** | Anti-rabbit | Sigma Aldrich | AB_1854720 | 1:300 |
| **PAX6** | Anti-rabbit | MBL | AB_1520876 | 1:500 |
| **PHH3** | Anti-rabbit | Sigma Aldrich | AB_477043 | 1:500 |
| **SMPD3** | Anti-rabbit | Abcam | n/a | 1:200 |
| **SMPD4** | Anti-rabbit | Sigma Aldrich | AB_2680762 | 1:200 |
| **Ceramide (C16)** | n/a | Avanti Polar Lipids | n/a | 2μM |
| **Fumonisin B1** | n/a | Sigma Aldrich | n/a | 30μM |
| **GW4869** | n/a | Tocris Bioscience | n/a | 10μM |

**Supplementary Table 4:**
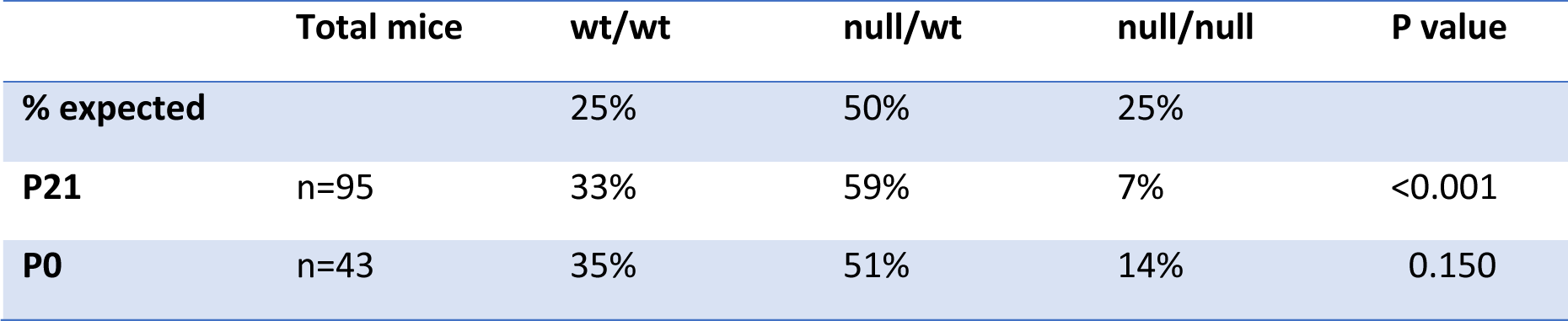
*Smpd4 null* survival.

**Supplementary Table 5:** *Smpd4 En1-Cre* conditional survival.

|  | Total mice | no Cre; flox/wt | En1-Cre+; flox/wt | no Cre; null/flox | En1-Cre+; null/flox | p value |
| --- | --- | --- | --- | --- | --- | --- |
| **% expected** |  | 25% | 25% | 25% | 25% |  |
| **P21** | n=38 | 13% | 24% | 16% | 13% | 0.178 |
| **P0** | n=46 | 33% | 20% | 20% | 26% | 0.695 |

**Supplementary Table 6:**
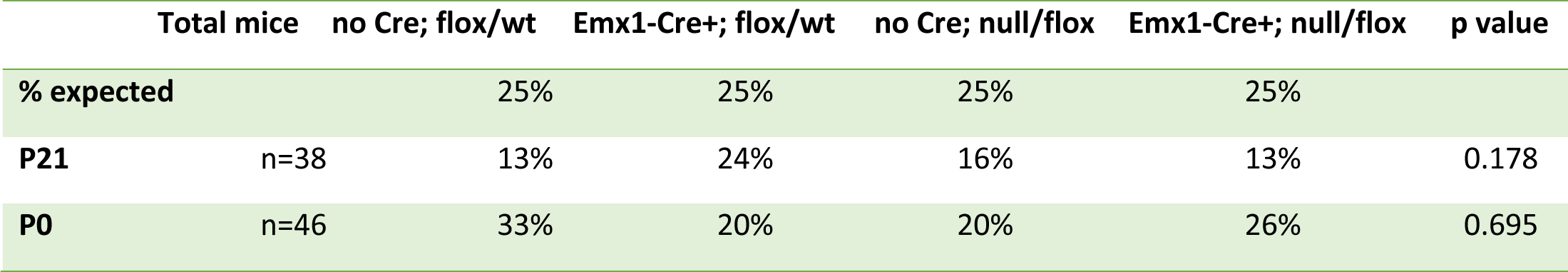
*Smpd4 Emx1-Cre* conditional survival.

**Supplementary Table 7:** *Smpd4 Foxg1-Cre* conditional survival.

|  | Total mice | no Cre; flox/wt | Foxg1-Cre+; flox/wt | no Cre; null/flox | Foxg1-Cre+; null/flox | p value |
| --- | --- | --- | --- | --- | --- | --- |
| **% expected** |  | 25% | 25% | 25% | 25% |  |
| **P21** | n=24 | 33% | 33% | 13% | 21% | 0.370 |

**Supplementary Table 8:** *Smpd3* deletion survival (C57B6J)

|  | Total mice | *Smpd3* wt | *Smpd3* del/wt | *Smpd3* del/del | p value |
| --- | --- | --- | --- | --- | --- |
| **% expected** |  | 25% | 50% | 25% |  |
| **P21** | n=33 | 39% | 61% | 0% | 0.003 |
| **P0** | n=35 | 40% | 60% | 0% | 0.002 |
| **E18.5** | n=46 | 30% | 59% | 11% | 0.086 |

**Supplementary Table 9:** *Smpd3* deletion survival (CD1)

|  | Total mice | *Smpd3* wt | *Smpd3* del/wt | *Smpd3* del/del | p value |
| --- | --- | --- | --- | --- | --- |
| **% expected** |  | 25% | 50% | 25% |  |
| **P0** | n=29 | 45% | 55% | 0% | 0.003 |

**Supplementary Table 10:**
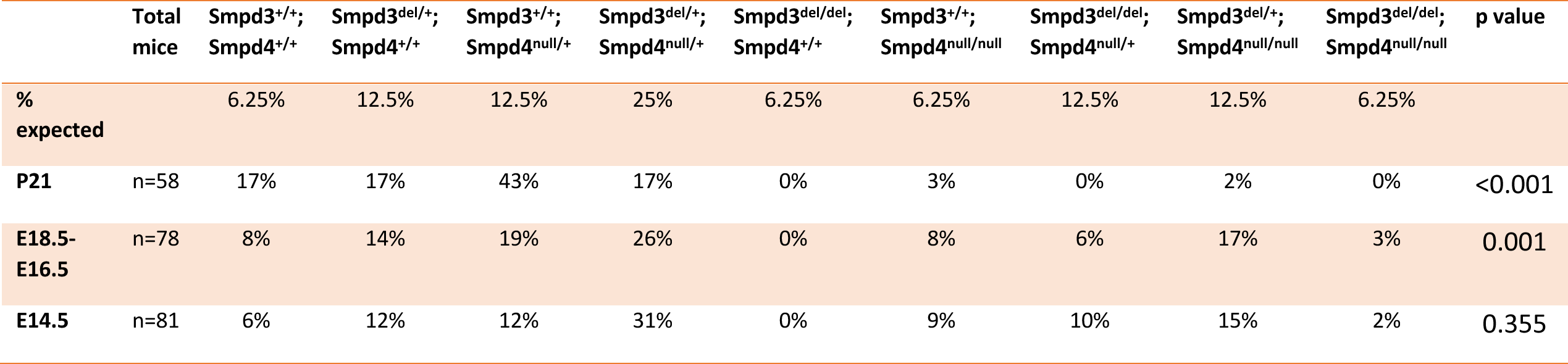
*Smpd3; Smpd4* double knockout survival.

## Notes

### Competing Interest Statement

The authors have declared no competing interest.

